# Comprehensive metabolic engineering for fermenting glycerol efficiently in *Saccharomyces cerevisiae*

**DOI:** 10.1101/2021.02.13.430370

**Authors:** Sadat M. R. Khattab, Takashi Watanabe

## Abstract

Glycerol is an eco-friendly solvent enhancing plant-biomass decomposition through a glycerolysis process in many pretreatment methods. Nonetheless, the lack of efficient conversion of glycerol by natural *Saccharomyces cerevisiae* restrains many of these scenarios. Here we outline the complete strategy for the generation of efficient glycerol fermenting yeast by rewriting the oxidation of cytosolic nicotinamide adenine dinucleotide (NADH) by O_2_-dependent dynamic shuttle while abolishing both glycerol phosphorylation and biosynthesis pathways. By following a vigorous glycerol oxidative pathway, the engineered strain demonstrated augmentation in conversion efficiency (CE) reach up to 0.49g-ethanol/g-glycerol—98% of theoretical conversion—with production rate >1 g/L^-1^h^-1^ when supplementing glycerol as a single fed-batch on a rich-medium. Furthermore, the engineered strain showed a new capability toward ferment a mixture of glycerol and glucose with producing >86 g/L of bioethanol with 92.8% of the CE. To our knowledge, this is the highest ever reported titer in this regard. Notably, this strategy flipped our ancestral yeast from non-growth on glycerol, on the minimal medium, to a fermenting strain with productivities 0.25-0.5 g/L^-1^h^-1^ and 84-78% of CE, respectively and 90% of total conversions to the products. The findings in metabolic engineering here may release the limitations of utilizing glycerol in several eco-friendly biorefinery approaches.

**IMPORTANCE:** With the avenues for achieving efficient lignocellulosic biorefinery scenarios, glycerol gained keen attention as an eco-friendly biomass-derived solvent for enhancing the dissociation of lignin and cell wall polysaccharides during pretreatment process. Co-fermentation of glycerol with the released sugars from biomass after the glycerolysis expands the resource for ethanol production and release from the burden of component separation. Titer productivities are one of the main obstacles for industrial applications of this process. Therefore, the generation of highly efficient glycerol fermenting yeast significantly promotes the applicability of the integrated biorefineries scenario. Besides, the glycerol is an important carbon resource for producing chemicals. Hence, the metabolic flux control of yeast from glycerol contributes to generation of cell factory producing chemicals from glycerol, promoting the association between biodiesel and bioethanol industries. Thus, this study will shed light on solving the problems of global warming and agricultural wastes, leading to establishment of the sustainable society.

## Introduction

One of the challenges for sustaining the future humanosphere is to produce required bio-based chemicals and fuels from renewable resources to reduce greenhouse gas emissions. A massive requirement for ethanol recently arose for use in sanitizers because of the COVID-19 pandemic. Bakers’ yeast (*Saccharomyces cerevisiae*) has several desirable characteristics during bioethanol production, such as the high producer with a long and safe history of use. Besides, its unicellular structure, short life cycle, remarkable tolerance to inhibitors and stress during the industrial processes. Hence has been selected as a top model platform of microbial cell factories for several biotechnological applications (1, 2).

The resources of fermentable sugars are limited. Therefore, challenges exist to overcome the drawbacks of production from lignocellulosic biomass through evolving the maximum efficiencies in ethanol production from xylose and acetic acid with glucose (3-5). In the past decade, glycerol-producing industries, particularly that of biodiesel, have expanded and accumulated substantial amounts of glycerol, which has led to price drops (6). Although the reduction degree of glycerol (C_3_H_8_O_3_) is higher than that of other fermentable sugars (7), glycerol is classifying as a non-fermentable carbon in native *S. cerevisiae* (2). Additionally, glycerol is a carbon source poorly utilized primarily via the glycerol-3-phosphate pathway (herein referred to as the G3P pathway), which is composed of glycerol kinase (*GUT1*) and FAD-dependent-mitochondrial glycerol-3-phosphate dehydrogenase (*GUT2*) (8). Yeast biosynthesis glycerol for mitigating osmotic stress and optimizing the redox balance (9). Furthermore, glycerol catabolism is subject to the repression and transcriptional regulation of glucose through respiratory factors and the *GUT1* and *GUT2* genes (10-12).

The trials to ferment glycerol in *S. cerevisiae* by overexpressing the native oxidative pathway (DHA; dihydroxyacetone pathway) have been reported. Overexpress endogenous glycerol dehydrogenase (*ScGCY1)* with dihydroxyacetone kinase (*ScDAK1*) produced 0.12 g ethanol/ g glycerol (g^e^/g^g^). The production rate was 0.025 g/L^-1^h^-1^ (13). The importance of glycerol as a carbon source that can be utilized by yeast cells has further prompted the study, which revealed the intraspecies diversity ranging from good glycerol grower to non-growers among 52 *S. cerevisiae* studied strains. The growth phenotype is controlled by quantitative traits (14). It has been reported that many alleles, such as cytoplasmic ubiquitin-protein ligase E3, *UBR2*, link *GUT1* for growth on glycerol in synthetic medium without supporting supplements (15). On the other hand, heterologous replacement of G3P with DHA combined with the glycerol facilitator (*FPS1*) resulted in the restoration of growth characteristics like the parental strain or even higher (16). This replacement in a positive-glycerol grower ancestor with limiting the oxygen in shaking flask cultures increased the ethanol production 0.165 to 0.344g^e^/g^g^ in buffered synthetic medium. The maximum titer produced reached 15.7 g/L after 144h (17). So far, glycerol is reporting as a non-fermentable carbon source in *S. cerevisiae*, and attempts are undergoing to ferment this carbon in this yeast (2). Meanwhile, the methylotrophic yeast *Ogataea polymorpha* has been tested for bioethanol production from glycerol by overexpressing multiple genes involved in either the DHA or G3P pathways. In addition to *FPS1*, pyruvate decarboxylase (*PDC*) 1, and alcohol dehydrogenase (*ADH*) 1. The overall ethanol production was 10.7 g/L ethanol as a maximum accumulated product at an efficiency of 0.132 g^e^/g^g^. (18). Hitherto, there is no known native or genetically engineered *S. cerevisiae* strain ferments glycerol efficiently to ethanol.

Of note, we previously reported microwave-assisted pretreatments of recalcitrant lignocellulosic biomass in aqueous glycerol (19) and further innovated that method with the catalysis of alum [AlK(SO4)2] (20). Fractionation of sugars from glycerol is a costly process. Therefore, to improve our scenario, there is a need for developing an *S. cerevisiae* capable of efficiently fermenting glycerol with glucose following glycerolysis of biomass. Such glycerol-fermenting strain will co-ferment glycerol to synergist ethanol production from hydrolyzed lignocellulosic biomass to an economical distillation process. In the present study, we report the details of modelling the yeast cell to redirect glycerol traffic to bioethanol production to th highest ever reported >8.6%—with the presence of glucose—through innovation of the forthcoming systematic metabolic engineering outlined in Figure 1 as follows: 1) abolishment of the inherent glycerol biosynthesis pathway by knocking out NAD-dependent glycerol-3-phosphate dehydrogenase (*GPD*) 1; 2) replacement of cytosolic NADH oxidation through the *ScGPD1* shuttle with a more effective O_2_-dependent dynamic shuttle of water forming the NADH oxidase *NoxE*; 3) knocking out the first gene of the G3P pathway (*GUT1*); and 4) implementing a vigorous oxidative pathway via overexpression of two copies of both the heterologous genes of glycerol dehydrogenase from *O. polymorpha* (*OpGDH*) and the glycerol facilitator *CuFPS1*, besides two copies of both the endogenous genes *ScTPI1* and *ScDAK1* with one copy of *ScDAK2*.

**Fig. 1.**
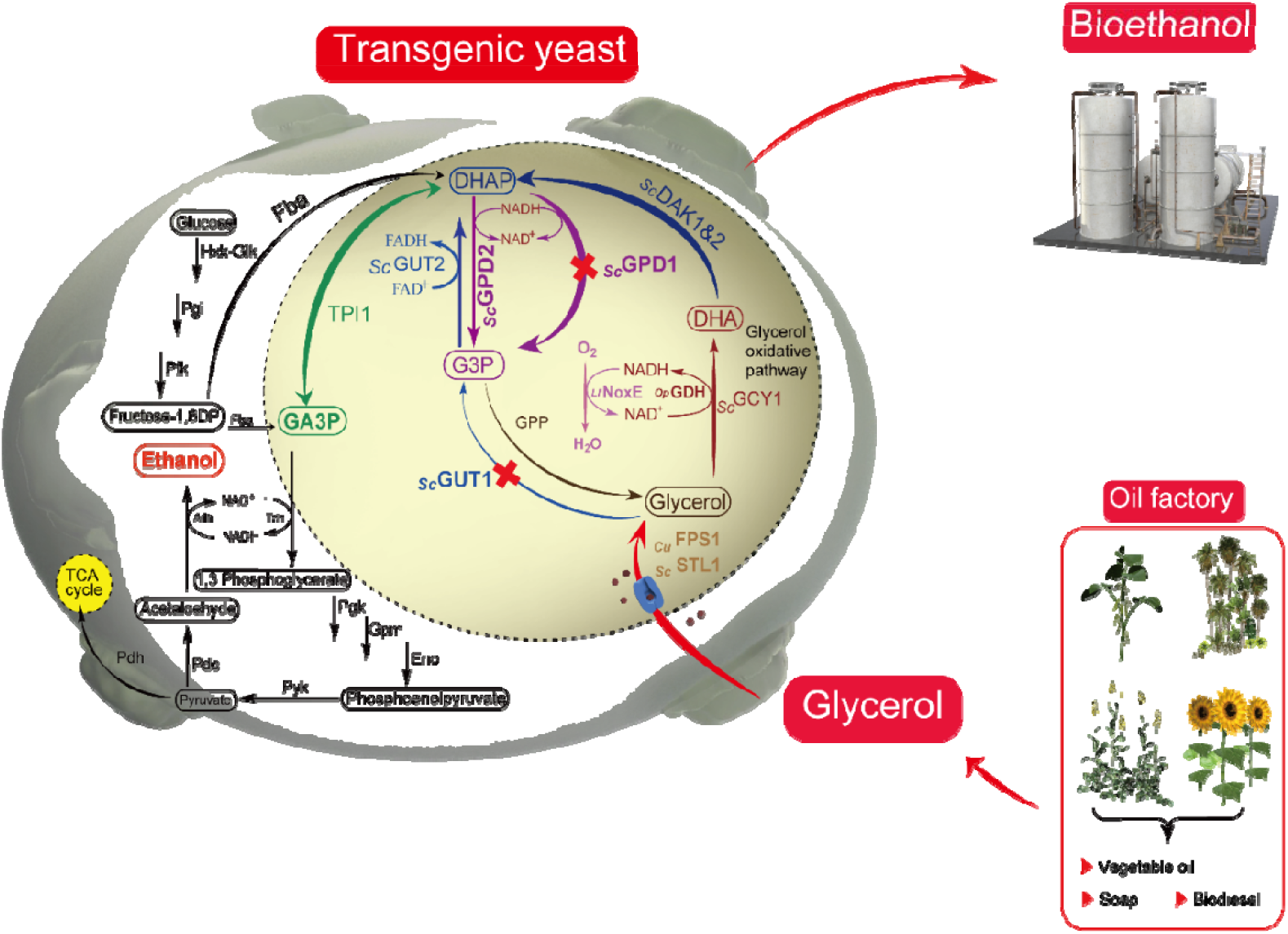
Schematic diagram showing the integrative scenario of a biorefinery with new generation of glycerol fermenting yeast and redirection of glycerol influxes to ethanol production in *Saccharomyces cerevisiae* via retrofitted native glycerol anabolic and catabolic pathways using the robust oxidative route with renovation of the NAD^+^ cofactor via O_2_-dependent dynamics of water-forming NADH oxidase. During pathway re-routing, glycerol-3-phosphate dehydrogenase 1 (*ScGPD1*) and glycerol kinase 1 (*GUT1*) were knocked-out. A highlighted circle indicates the overexpressed indigenous *S. cerevisiae* enzymes dihydroxyacetone kinase (*ScDAK*) 1 and 2 as well as triosephosphate isomerase (*ScTPI*) 1, heterologous glycerol dehydrogenase from *Ogataea polymorpha* (*OpGDH*), glycerol facilitator from *Candida utilis* (*CuFPS1*) and water-forming NADH oxidase from *Lactococcus lactis* subsp. *lactis* Il1403 (*LlNoxE*).

## MATERIAL AND METHODS

### Strains, primers, cassettes, and plasmids constructions

Ancestor and all recombinant strains used in this study are listed in Table 1. The original ancestors showed in Fig. S1. The plasmids used in this study are listed in Table 2. The primers used here are listed in Table S1, and those for obtaining the native genes were designed based on the sequences available on the S288C *Saccharomyces* Genome Database (SGD): https://www.yeastgenome.org/(S288C). Details of DNA fragments, cassettes, and plasmids construction are explained as follows:

**Table 1.** Characteristics of *S. cerevisiae* strains generated in this study:

| *S. cerevisiae* strain | Relevant genotype | Reference |
| --- | --- | --- |
| D452-2 | MATa *leu2 his3 ura3 can1* | (25) |
| D452-2-URA3 | D452-2, URA3:: TDH3 promoter and d22DIT1 terminator | This study |
| GDH | D452-2, URA3:: TDH3p-*OpGDH*-d22-DIT1t | This study |
| GPD1/LINoxE (NOXE) | D452-2, Δ*GPD1*:: TDH3p-*LINoxE*-d22-DIT1t | this study |
| GDH-NOXE (GN) | D452-2, URA3:: TDH3p-*OpGDH*-d22-DIT1t; Δ*GPD1*:: TDH3p-*LINoxE*-d22-DIT1t | This study |
| D452-2-URA3-AUR1-C | D452-2, URA3:: TDH3 promoter and d22DIT1 terminator :: AUR1-C | This study |
| GN-FDT | D452-2, URA3:: TDH3p-*OpGDH*-d22-DIT1t; Δ*GPD1*:: TDH3p-*LINoxE*-d22-DIT1t; AUR1-C:: PGKp-*CuFPS1*-RPL41Bt; PGKp-*ScTPII*-PGKt; PGKp-*ScDAK2*-PGKt; PGKp-*ScDAK1*-PGKt | This study |
| GN-FDT-M1 (SK-FGG) | D452-2, URA3:: TDH3p-*OpGDH*-d22-DIT1t; Δ*GPD1*:: TDH3p-*LINoxE*-d22-DIT1t; AUR1-C:: PGKp-*CuFPS1*-RPL41Bt; PGKp-*ScTPII*-PGKt; PGKp-*ScDAK2*-PGKt; PGKp-*ScDAK1*-PGKt; Δ*GUT1*:: TEFp-*CuFPS1*-CYC1t; TYS1p-*OpGDH*-ATP15t; TDH3p-*ScDAK1*-d22-DIT1t; FBA1p-*ScTPII*-TDH3t | This study |

**Table 2:** Constructed plasmids in this study.

| Plasmid | Relevant genotype | Reference |
| --- | --- | --- |
| pPGK-URA3 | URA3; PGK promoter and terminator | (21) |
| pPGK-DAK1 | URA3; expresses DAK1 | This study |
| pPGK-DAK2 | URA3; expresses DAK2 | This study |
| pPGK-TPI1 | URA3; expresses TPI1 | This study |
| pPGK-TPI1, *Sc*DAK2 | URA3; expresses TPI1, DAK2 | This study |
| pPGK-TPI1, *Sc*DAK2, *Sc*DAK1 | URA3; expresses TPI1, DAK2, DAK1 | This study |
| TDH3p-d22DIT1t- URA3 | URA3, TDH3 promoter and d22DIT1 terminator | This study |
| TDH3-*Op*GDH-d22DIT1t-URA3 | URA3; expresses *Op*GDH | This study |
| TDH3-LINoxE-d22 DIT1t-URA3 | URA3; expresses *L*INoxE | This study |
| pAUR101 | AUR1-c | Takara Bio |
| pAUR101-PGKp-RPL41Bt | AUR1-C, PGKp-RPL41Bt | This study |
| pAUR101-PGKp-*Cu*FPS1-RPL41Bt | AUR1-C; expresses *Cu*FPS1 | This study |
| pAUR101-*Cu*FPS1, TPI1, DAK2, DAK1 | AUR1-C; expresses *Cu*FPS1, TPI1, DAK2, DAK1 | This study |
| pAUR101- M1* | AUR1-C; expresses *Cu*FPS1, *Op*GDH, DAK1, TPI1 | This study |
| Multiplex pCAS (Addgene #60847) | Multiplex expresses Cas9 and HDV ribozyme-sgRNA | (23) |
| Multiplex pCAS/GPD1-1 | Multiplex expresses Cas9 and gRNA to base No. 135 of *GPD1* | This study |
| Multiplex pCAS/GPD1-2 | Multiplex expresses Cas9 and gRNA to base No. 1045 of *GPD1* | This study |
| Multiplex pCAS/GUT1 | Multiplex expresses Cas9 and gRNA to base No. 616 of GUT1 | This study |

\*M1: TEFp-CuFPS1-CYC1t; TYS1p-OpGDH-ATP15t; TDH3p-ScDAK1-d22-DIT1t; FBA1p-ScTPI1-TDH3t

### Construct pPGK-ScTPI1, ScDAK2, ScDAK1 plasmids

We obtained the genes from genomic DNA of the ancestor strain D452-2 to clone the plasmids in this section. Initially, cell walls were disrupted by resuspension with picked cells in 20 µl of 30 mM NaOH at 95°C for 10 min, and then used directly as a template for PCR; 1µl of freshly disrupted cells are suitable for a 50 µl of PCR mixture. High fidelity polymerisation of KOD-plus neo with their corresponded primers (Table S1, section 1) was used during this amplification. The Xhol site of *DAK2* was deleted before cloning. Genes were purified from the PCR mixtures FastGene Gel/PCR Extraction Kit (Nippon Genetics Co. Ltd), and their cohesive ends were formed according to the designated primers and restriction enzymes. We first separately cloned each gene in the pPGK/URA3 plasmid (21) under the control of the expression system PGK promoter and PGK terminator (Table 2). We obtained the plasmids and confirmed their gene sequences via sequencing using primers (Table S1, section 2). Next, we cut the XhoI/SalI-ScDAK2 cassette and inserted it into the SalI site of the pPGK-ScTPI1 plasmid. The non-reopened ligations (XhoI/SalI sites) were used during the ligation of new cassettes of DAK1 to form the new plasmid (Table 2).

### Construct TDH3p-d22DIT1t, TDH3-d22-OpGDH and TDH3-d22-LlNoxE plasmids

We first purchased the mutated terminator d22DIT1t from Integrated DNA Technology (IDT; Tokyo, Japan) according to the published sequences (22). Flanking sequences downstream of *ScGPD1* were added by the feature of PCR polymerization, with primers possessing a long tail to form fragment 2. TDH3 promoter magnified from the genomic DNA of the ancestor strain D452-2 with flanking sequences upstream of the *ScGPD1* by PCR to form fragment 1. A further extension to those flanking sequences may be accomplished by PCR when needed (Table S1, section 3). Then, cohesive ends of those coupled DNA fragments were processed by the restriction enzymes XhoI, NotI for the first fragment, and NotI, SalI for the second. After purification of the fragments, one-step cloning was employed coupled the TDH3 promoter and mutated DITI terminator into XhoI/SalI of PGK/URA3 plasmid. Then the TDH3p-d22DIT1t-URA3 plasmid was constructed (Table 2). Subsequently, we cloned the *OpGDH* gene into the constructed TDH3-d22-DIT1t plasmid to form the TDH3-OpGDH-d22DIT1t-plasmid. Synthetic *OpGDH* was also purchased from IDT according to the sequence deposited in GenBank under the accession number XP_018210953.1. Full sequences are available in Table S2. Also, we bought the water-forming NADH oxidase gene of *Lactococcus lactis* subsp. *lactis* Il1403 (IDT) based on the sequence on gene bank accession number AAK04489.1. Then cloned it into TDH3p-d22DIT1t to assemble TDH3-d22-LlNoxE plasmid (Tables 2).

### Construct multiplex pCAS-gRNA-CRISPR systems

The multiplex pCAS-gRNA system was a gift from Prof. Jamie Cate (Addgene plasmid # 60847; https://www.addgene.org/60847/) (23). We used an online tool for the rational design of CRISPR/Cas target to allocate the highest probability of the on-target sites for the gRNA in the genomic DNA of *S. cerevisiae* (https://crispr.dbcls.jp/) (24). Accordingly, the primers were designed based on the previously allocated sequence (20 bp before the PAM), with another 20 bp from sgRNA or HDV ribozyme for overlap (Table S2, sections 3.2, and 6.1). First, PCR was used to synthesize two fragments from the template, the pCAS-gRNA plasmid. The first fragment was amplified using the forward primer called pCAS F—located upstream of the gRNA scaffold at the SmaI site of the multiplex plasmid—and antisense primer with a reverse sequence of the target gRNA. The second fragment was amplified by a forward primer, which has a sense sequence of gRNA, and the reverse primer called pCAS R—located downstream of the gRNA scaffold (Table S1, section 3.2 and 6.1). After purifying each DNA segment, overlapping and integration were carried out by PCR using the pCAS F and R primers. The produced fragment was then end-repaired to the SmaI-PstI sites for cloning into a truncated pCAS-gRNA plasmid with SmaI-PstI. As a result, a new multiplex pCAS-gRNA plasmid was formed. Steps have been repeatedly performed with constructing multiplex pCAS-gRNA plasmids targeting *ScGPD1, GUT1* genes (Table 2). We confirmed the newly constructed systems by sequencing their entire scaffolds.

### Construct pAUR101-CuFPS1 and pAUR101-CuFPS1, ScTPI1, ScDAK2, ScDAK1plasmids

*Candida utilis* (NBRC 0988) was obtained from the National Biological Resource Center (NBRC) of the National Institute of Technology and Evaluation (Japan) and was used as a template to get the gene glycerol facilitator *FPS1* (*CuFPS1*). The sequence of *CuFPS1* was included in the deposited gene bank accession number BAEL01000108.1. The original pAUR101 plasmid was purchased from Takara Bio, Inc., Japan, and the primers were used to establish this plasmid listed (Table S1, section 4). A full sequence for the cassette PGK-CuFPS1-RPL41Bt was showed (Table S2). First, we constructed a pAUR101-PGKp-RPL41Bt vector by one-step cloning of the SmaI-Not1 PGK promoter (fragment F1) and NotI-SacI-RPL41B terminator (F2) into the SmaI-SacI pAUR101 vector and then cloning a cohesive ended NotI-*CuFPS* gene into the dephosphorylated NotI site of pAUR101-PGK-RPL41B vector to assemble pAUR101-PGKp-*CuFPS1*-RPL41Bt vector. We confirmed the clone direction by PCR. To constitute the pAUR101-CuFPS1, ScTPI1, ScDAK2, ScDAK1 plasmid, we detached the set of cassettes—ScTPI1, ScDAK2, and ScDAK1—from the previously constructed plasmid, pPGK-ScTPI1, ScDAK2, and ScDAK1 (Table 2) using restriction enzymes Xhol-SalI and re-inserted that set of cassettes into the SalI site of the pAUR101-PGK-CuFPS1-RPL41B plasmid (Table 2).

### Construct Module M1 and pAUR101-M1 plasmid

We first obtained all fragments which could form the module M1 separately by PCR (Fig. 2); the *CuFPS1, OpGDH* genes, and mutated d22DIT terminator amplified from their synthetic DNA stocks, whereas other fragments were magnified from the genomic DNA of the D452-2 strain (25). The full sequence of the module M1 is also accessible (Table S2), and the primer details are listed in Table S1, Section 5. Purification of the 12 amplified DNA pieces was carried out on 1%–2% agarose gel and then recovered by the extraction Kit. We accordingly obtained highly purified fragments before the onset of assembly using the Gibson Assembly Master Mix. We first joined every three consecutive segments seamlessly according to the manufacturer’s protocol (Gibson). We also directly amplified each set by PCR and then purified these again on an agarose gel. We repeatedly gathered every six sequential segments and then assembled the whole module M1; TEFp-CuFPS1-CYC1t; TYS1p-OpGDH-ATP15t; TDH3p-ScDAK1-d22-DIT1t; FBA1p-ScTPI1-TDH3t (Fig.2). We further added the SacI site upstream of the module M1 and the SmaI site downstream. These restriction sites were provided for cloning the module M1 into SacI-SmaI sites of pAUR101 vector to form pAUR 101-M1 (Table 2). Finally, we transferred the vector pAUR 101-M1 into *E. coli* as described previously and confirmed the accurate structure of M1 by sequencing the whole module M1 from pAUR-M1.

**Fig. 2.**
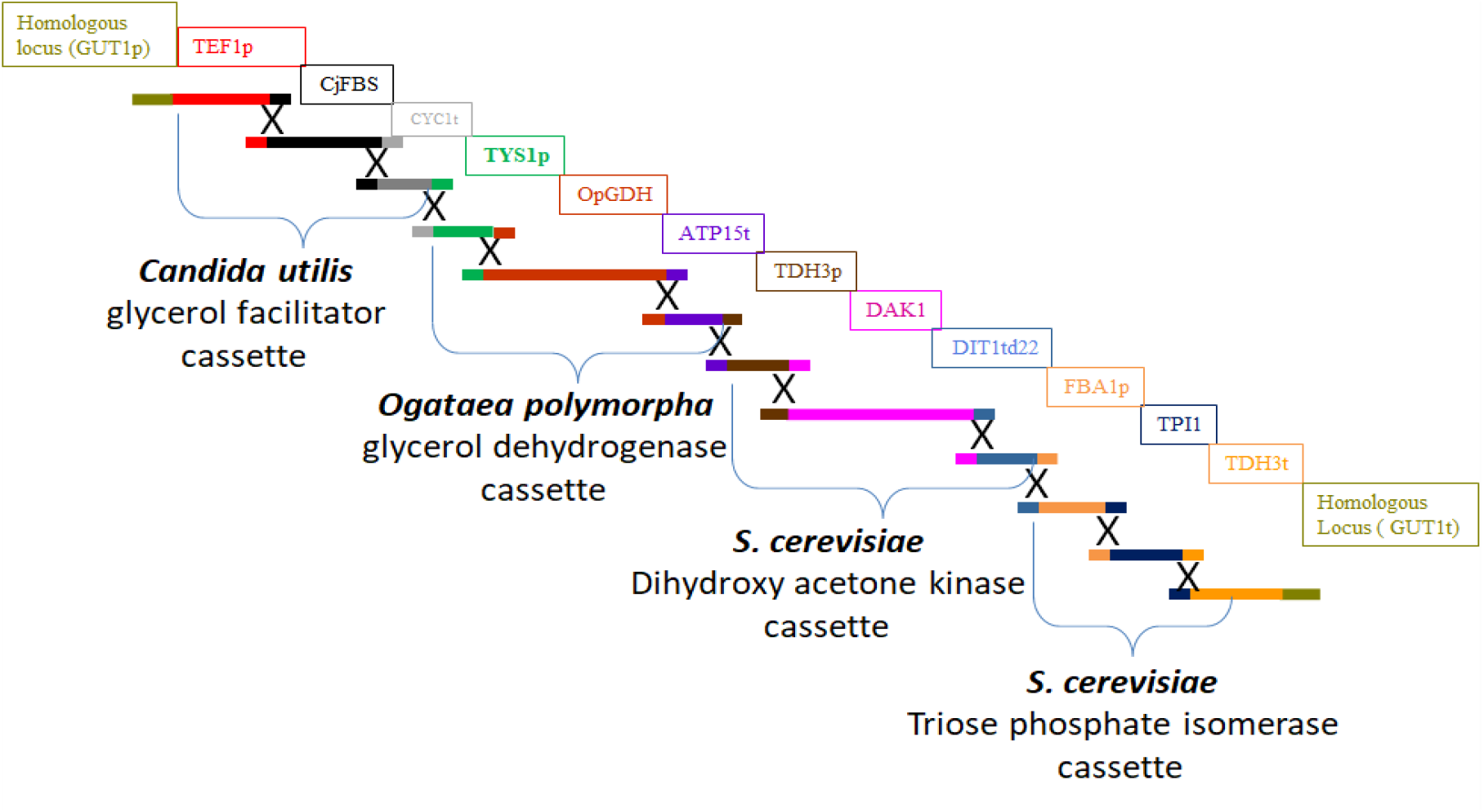
Depicted module M1 for replacing *ScGUT1* by homologous recombination using Multiplex CRISPR Cas 9.

### Transformation and recombination of strains

All the previous plasmids were stored in *E. coli* NEB 10-beta for further use, where the heat shock method was used during the transformation according to the procedures provided with the competent cells. All plasmid extractions were performed using the QIAprep Spin Miniprep Kit following the manufacturer’s protocol. All measurement of DNA was estimated using BioSpec-nano (Shimadzu, Japan). DNA was stored at −20°C for future use. Yeast transformation was carried out by Fast Yeast Transformation ™kit (Takara Bio) to integrate linear pAUR101 vector and its associated genes in the AUR1-C locus. Also, linear pPGK plasmid with its cloned genes in URA3 loci (25). For achieving genome editing and the replacement of *ScGPD1*and *ScGUT1* genes with its designated DNA repairing cassette or module, we used the protocol of CRISPR-Cas9 genome engineering in *S. cerevisiae* cells (26). We confirmed target replacements using PCR for the inserted repairing cassettes with primers from upstream and downstream of the flanking recombined loci. We re-confirmed the recombination after the losses of pCAS multiplex plasmid. All recombination strains and their genotypes are listed in Table 1.

### Preparation of cell-free extract

Cell proteins were extracted as previously described (3, 27) with some modifications. The recombinant strains listed in Table 1 were cultivated in closed 50-ml Falcon tubes with 10 ml of YP medium supplemented with (w/v) 1.5% glucose and 7% glycerol for 15h. Cell pellets were harvested by centrifugation at 700 xg for 2 min at 4 °C, then washed with 20 ml of 100 mM of HEPES buffer (pH 7.4) and centrifuged again. Then, the cell pellets were lysed in 1 ml of HEPES buffer supplemented with 1 mM MgCl_2_ and 10 mM 2-mercaptoethanol with approximately 400 mg of glass beads in a 2-ml Eppendorf tube. The lysis was accomplished by vigorous shaking by bench vortex with six-time intervals on ice for 30 seconds. The crude proteins were separated from the glass beads and cell debris by two rounds of centrifugation at 22300 xg at 4°C for 5 min. Total protein concentration was estimated using Bio-Red, Quick Start ™Bradford 1 x dye reagent by measuring the absorbance at 595nm using BSA as a standard.

### Enzyme assays

The specific activity of glycerol dehydrogenase was assayed by monitoring the increases of NADH absorbance at 340 nm. UV-spectrophotometer measured the slope of change (UV-2700; UV-VIS spectrophotometer, Shimadzu, Japan). One ml mixture of 80 mM HEPES buffer (pH 7.4), 5 mM NAD^+^,100 mM glycerol, and 10 µl of crude extract were mixed during analysis. The activity of *ScTPI1* was assayed as described previously (28) with some modifications. The reaction mixture was 1 ml in volume and was composed of 100 mM triethanolamine hydrochloride (pH 7.53), 2.5 mM NAD^+^, 10 mM DHAP, and 10 µl of crude extract. Dihydroxyacetone kinase was assayed using a universal kinase activity kit according to the manufacturer’s instructions (Catalog Number EA004).

### Intracellular concentration of NAD^+^ /NADH

The cellular contents of NAD^+^/NADH were colorimetric quantitatively determined at 565 nm using a BioAssay Systems (E2ND-100) EnzyChrom™NAD^+^ /NADH Assay Kit.

### Fermentation procedures

The fermentation experiments were performed in several conditions defined as following according to the ratio of liquid medium (ml): Erlenmeyer flasks volume (ml). Micro-aerobic [20:100], semi-aerobic [20:200], aerobic [20:300]. We conducted the fermentation at 30 ± 2 °C with agitation speeds at 180 or 200 rpm. For strict anaerobic conditions, 20 ml of YNB medium with initial cell pellets were transferred into a sterilized 50-ml vial and then capped with a precision seal septa cap. Afterward, the nitrogen gas was flushed into the culture medium through Terumo needles. Sampling was drawn through needles. The fermenter cells were harvested from the same volume of the pre-culture YPD medium for approximately 15h. For SK-FGG, the cultivation culture medium was YPD_20_G_70_. Cells were harvested by centrifugation at 1600 ×g for 5 min at 4°C and washed with sterile water. Then, collected cells were re-supplemented with the YP medium with glucose, glycerol, or both, as shown in figures and tables. These concentrations have been chosen to match those can obtain after the glycerolysis of biomass (20). The cell density was monitored using spectrophotometry at 600 nm (AS ONE, Japan).

### Fermentation analysis

All analyses were performed using ultra-fast liquid chromatography (Shimadzu, Japan). Automatic sampling was fractionated by Aminex HPX-87H column (Bio-Rad Laboratories, Hercules, CA, USA), then analyzed in a refractive index detector (RID-10A; Shimadzu) and a prominence diode array detector (SPD-M20A; Shimadzu). Fractionation was accomplished at a flow rate of 0.6 ml/min with 5 mM H_2_SO_4_ as the mobile phase at 50°C. Reactant concentrations were estimated by monitoring the peak areas compared with the standards of the authenticating reactant’s glucose, glycerol, ethanol, acetic acid, pyruvate, succinic acid, and acetaldehyde, which detected by RID. We used the SPD to estimate DHAP and DHA. The detectable quantities were reported in the tables or figures and were omitted if it was undetectable.

## RESULTS

### Overexpress of a vigorous glycerol dehydrogenase

The overexpression of *OpGDH* in the GDH strain was verified by comparing the enzyme activity with the reference D452-2 (Table 3). As a result, 58% of glycerol was consumed compared with 8% from reference strain, which raised ethanol level from 4.47 in ancestral to 11.82g/L in GDH strain, representing 0.27g^e^/g^s^ before reutilizing the produced ethanol after 26h (Fig. 3).

**Table 3.** Specific activities of the enzymes; glycerol dehydrogenase (*GDH*), dihydroxyacetone kinase (*DAK*), triosephosphate isomerase (*TPI1*) and NADH/NAD^+^ ratio with their intracellular concentrations in the recombinant strains in this study. Error values represent standard deviation of the mean (n = 2). * indicates not detected. ¤ indicates not measured.

| Relevant<br>Strain | Enzyme activities $\mu\text{mole}/\text{min}/\text{mg}$ cell<br>extracted proteins | | | Intracellular concentrations | | NADH/ NAD<sup>+</sup><br>Ratio |
| --- | --- | --- | --- | --- | --- | --- |
| | *GDH* | *DAKs* | *TPII* | NADH($\mu\text{M}$) | NAD<sup>+</sup> ($\mu\text{M}$) | |
| WT | ND* | $3.9 \pm 0.23$ | $0.058 \pm 0.003$ | $0.17 \pm 0.004$ | $0.36 \pm 0.01$ | $0.47 \pm 0.01$ |
| GDH | $0.063 \pm 0.001$ | NM∅ | $0.055 \pm 0.003$ | $0.26 \pm 0.008$ | $0.38 \pm 0.02$ | $0.68 \pm 0.02$ |
| NOXE | ND* | NM∅ | $0.054 \pm 0.002$ | $0.04 \pm 0.005$ | $0.36 \pm 0.04$ | $0.12 \pm 0.01$ |
| GN | $0.065 \pm 0.004$ | NM∅ | $0.057 \pm 0.003$ | $0.34 \pm 0.018$ | $0.35 \pm 0.01$ | $0.99 \pm 0.02$ |
| GN-FDT | $0.071 \pm 0.003$ | $7.15 \pm 0.23$ | $0.067 \pm 0.004$ | $0.17 \pm 0.08$ | $0.52 \pm 0.01$ | $0.33 \pm 0.02$ |
| SK-FGG | $0.111 \pm 0.002$ | $18.3 \pm 0.28$ | $0.076 \pm 0.004$ | $0.11 \pm 0.01$ | $0.63 \pm 0.01$ | $0.18 \pm 0.01$ |

**Fig. 3.**
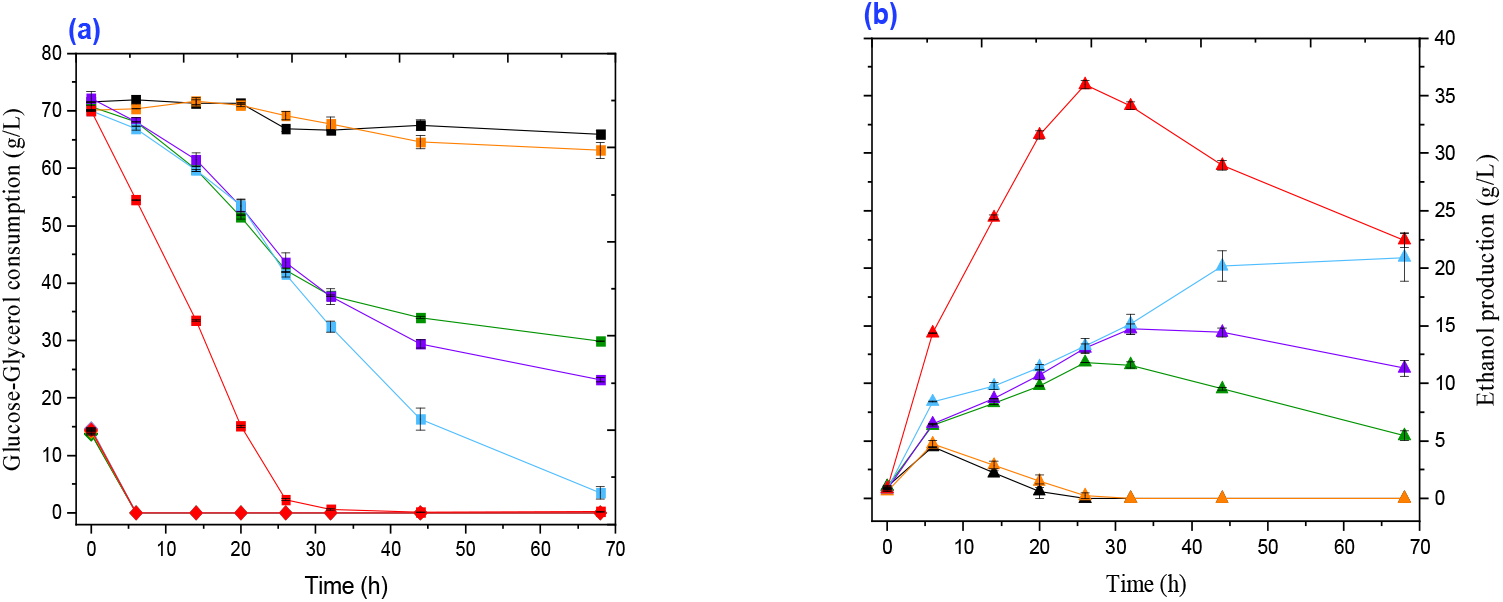
Comparison of the time course of glycerol–glucose fermentation between the *S. cerevisiae* strains within this study: Reference ancestral strain (black lines); NOXE strain (orange lines); GDH strain (green lines); GN strain (purple lines); GN-FDT strain (blue lines); GN-FDT-M1[SK-FGG] (red lines); (a) Glucose consumption (rhomboid symbols), glycerol consumption (squares); (b) ethanol production (triangles). Fermentation carried out in semi-aerobic conditions in flasks with 20:200 medium: flask volume at 30°C with shaking at 180 rpm. YP medium supplemented with (w/v) 1.5% glucose, and 7% glycerol was used. (n= 3).

### Integrate the replacement of *ScGPD1* by *LlNoxE* with *OpGDH*

Based on data from the comparative study of substituting GPD shuttles with *LlNoxE*, the replacement of *ScGPD1* by *LlNoxE* abolished glycerol biosynthesis by 98% (Khattab and Watanabe, manuscript in preparation). Therefore, we studied the recycling outputs of NAD^+^/NADH between *OpGDH* and *LlNoxE* genes. We projected abolishing glycerol formation and decreases the ramification of DHAP, which consolidates straightforward progression to the glycolysis route and ethanol formation. This innovative integration clearly showed improvements in glycerol conversion efficiency to ethanol and delay in cell shift to utilize the produced ethanol. The ethanol production boosted to 13.27 g/L (0.31g^e^/ g substrate consumed) and extending the fermentation time to 32h with further raising the ethanol titer to 14.42 g/L. (Fig. 3). Analysis of a cofactor ratio (NADH/ NAD^+^) showed a significant change from 0.68 to 0.99 between GDH and GN strains, respectively (Table.3). The higher NADH/NAD^+^ ratio is pointing to the higher efficiency of replacing *ScGPD1* with *LlNoxE* shuttle.

### Overexpress other DHA pathway genes: *ScTPI1, ScDAK1, ScDAK2*, and *CuFPS1*

With current outputs, further limitations in the activity of other genes in the DHA pathway— *ScTPI1, ScDAK*s, and *CuFPS1*—affecting the traffic of glycerol conversion to ethanol. Therefore, we overexpressed all of them. In this point, we clarify that *ScDAK2* and *ScTPI1* are not considered during the previous studies of converting glycerol to ethanol. In the GN-FDT strain, the specific enzyme activity of *OpGDH* increased by 9% compared with the GN strain. Also, *ScDAKs* and *ScTPI1* increased by 83% and 16%, respectively, compared with the ancestor strain. Moreover, the NADH/NAD^+^ ratio decreased to 0.33% of the GN (Table 3). This recombination step (GN-FDT strain) unequivocally solved one of the problems in this study, where it prevented the shift in utilizing ethanol before the consumption of glycerol. The consumption rate reached 1 g/L^-1^h^-1^ and produced 20.95 g/L of ethanol using this recombinant strain. Nonetheless, the conversion efficiency appeared to be less than 48% of its theoretical value (Fig. 3).

### Reinforcement the DHA pathway using a further copy of *ScTPI1, ScDAK1, OpGDH*, and *CuFPS1* genes while abolishing the native G3P route

Boost the efficiency of glycerol fermentation by another copy of the DHA pathway was also studied. Accordingly, M1 was constructed by employing Gibson hybrid assembly and PCR to replace *ScGUT1* (Fig. 2). As a result of integration with replacement, the specific enzyme activities of *OpGDH, ScDAK*, and *ScTPI1* were further enhanced by 56%, 256%, and 13%, respectively, compared with the GN-FDT strain. Besides, the NADH/NAD^+^ ratio decreased to 0.11 (Table 3). Interestingly, we obtained outstanding fermentation in this fourth step of recombination GN-FDT-M1, named SK-FGG strain (Table 1, Fig. 3). The glycerol consumption rate reached 2.6 g/L^-1^h^-1^ at the described experimental conditions, and the productivity paced 1.38 g/L^-1^h^-1^ of ethanol with a conversion efficiency of 0.44g^e^/g^s^ (Fig. 3).

### Ferment powers in the engineered yeast

With the current state of metabolic engineering, we examined fermentation characteristics at higher initial concentrations of glycerol (110 g/L) in the absence and presence of glucose (22.5 g/L), where glucose was reported as a suppressor for glycerol fermentation (Table 4 [A and B]). We also tested fermenting a further fed-batching of 100 g/L glycerol to the condition of [B] in (Table 4 [C]). To further confirming the engineered strain is unsubjected to repression by glucose during the glycerol fermentation, we lifted the glucose level to 45 g/L with decreased the glycerol concentration by 25% to 82 g/L before further adding 100 g/L glycerol as a fed-batch (Table 4 [D]). These outlines also testing the strain SK-FGG to produce an economically distillable ethanol titer (Table 4 [D]). The concentrations of glucose and glycerol in [D] are relatively like those obtained after the glycerol process (20). The strain SK-FGG exhibited outstanding performance in semi-aerobic conditions at a higher initial concentration of glycerol–glucose mixture. Its conversion efficiency reached 98% (0.49 g^e^/g^g^) with a production rate of >1 g/L^-1^h^-1^ of ethanol after consumed 82.5 g/L of glycerol from one fed-batch condition (Table 4, condition [A]). Acetic acid accumulated at 1.14 g/L at this condition (Table 4, condition [A]). Even at mixing glycerol with 22.55 g/L of glucose, its conversion efficiency was comparatively the same (Table 4, condition [B]). Interestingly, the strain engineered here is exceptional in its capacity to harmonize fermenting glycerol with glucose, along with an accumulation of >86 g/L of bioethanol with additional fed batching of glycerol (Table 4, [D]). With higher initial glucose at condition [D], cell density was promoted by 31% compared with the case [C]; besides, a minor reduction of the efficiency of ethanol conversion (Table 4, [C and D]).

**Table 4.** Fermentation characteristics by the generated strain SK-FGG at high concentrations of glycerol, with and without adding glucose.

| Parameter | Fermentation conditions and products values |  |  |  |
| --- | --- | --- | --- | --- |
|  | One fed-batch | One fed-batch | Two fed-batch | Two fed-batch |
|  | [A] | [B] | [C] | [D] |
| Fermentation time (h) | 40 ± 0.30 | 40 ± 0.30 | 96 ± 0.30 | 96 ± 0.30 |
| Total consumed glycerol (g/L) | 82.5 ± 1.63 | 75.77 ± 1.34 | 141.88 ± 2.66 | 141.703 ± 4.79 |
| Total consumed glucose (g/L) | NA | 22.55 ± 1.14 | 22.2 ± 0.67 | 44.67 ± 0.9 |
| Rate of consumption (g/L<sup>-1</sup>h<sup>-1</sup>) | 2.06 ± 0.017 | 2.42 ± 0.014 | 1.71 ± 0.011 | 1.94 ± 0.015 |
| Ethanol yield (g/L) | 40.42 ± 0.89 | 48.22 ± 1.16 | 79.74 ± 1.03 | 86.54 ± 2.48 |
| Rate of ethanol production (g/L<sup>-1</sup>h<sup>-1</sup>) | 1.01 ± 0.01 | 1.2 ± 0.02 | 0.83 ± 0.01 | 0.9 ± 0.02 |
| Efficiency of ethanol production (g<sup>e</sup>/ g<sup>s</sup>) * | 0.49 ± 0.002 | 0.49 ± 0.002 | 0.486 ± 0.003 | 0.464 ± 0.003 |
| Efficient/ theoretical (%) <sup>+</sup> | 98 | 98 | 97.2 | 92.8 |
| Acetic acid accumulation (g/L) | 1.14 ± 0.02 | 1.12 ± 0.02 | 1.67 ± 0.1 | 2.46 ± 0.02 |
| Total conversion (g<sup>p</sup>/ g<sup>s</sup>) <sup>++</sup> | 0.5 ± 0.002 | 0.5 ± 0.003 | 0.496 ± 0.005 | 0.478 ± 0.004 |
| Cell Density (OD<sub>600</sub>) | 10.6 ± 0.2 | 11.6 ± 0.2 | 11.6 ± 0.3 | 15.2 ± 0.3 |

Oxygen availability in (liquid medium: flask volume) at 30°C with shaking at 200 rpm (20:200).
(g<sup>e</sup>/ g<sup>s</sup>) \*; gram ethanol per gram of glycerol. (g<sup>p</sup>/ g<sup>s</sup>) <sup>++</sup>; gram products per gram substrate
Efficient/ theoretical (%) <sup>+</sup>; C<sub>6</sub>H<sub>12</sub>O<sub>6</sub> → 2 C<sub>2</sub>H<sub>5</sub>OH + 2 CO<sub>2</sub> → **Eq. 1** C<sub>3</sub>H<sub>8</sub>O<sub>3</sub> + ½ O<sub>2</sub> → C<sub>2</sub>H<sub>5</sub>OH + CO<sub>2</sub> + H<sub>2</sub>O → **Eq. 2**
(1 g) → 0.51 g + 0.49 g (1 g) → 0.5 g + 0.478 g

### Ferment glycerol as a sole carbon source

Eventually, verifying the fermentation capability of SK-FGG has been conducted on glycerol as a sole carbon in a yeast nitrogen base (YNB) medium with only 20 mg/L of leucine and histidine in a single-batch. We compared the fermentation at four degrees of oxygen availabilities. Under the strict anaerobic conditions, there was no further growth nor production of ethanol (Table 5). Under micro-aerobic conditions, the strain consumed 37.17 g glycerol with a consumption rate that reached 0.62 g/L^-1^h^-1^ and a production rate of 0.25 g/L^-1^h^-1^. The efficiency of ethanol conversion approached 0.42 g^e^/g^g^. Acetic acid accumulated at 0.78 g/L in this condition (Table 5). Glycerol was consumed more rapidly in a semi-aerobic atmosphere. The consumption rate was >1 g/L^-1^h^-1^, which raised the ethanol production rate to 0.44 g/L^-1^h^-1^ till accumulate 20.97 g/L. There was 2.88 g of acetic acid cumulation in this condition. As a result, the total conversion reached 0.44 g/g glycerol (Table 5). In aerobic conditions, glycerol consumption and ethanol production were boosted to 1.29 and 0.5 g/L^-1^h^-1^, respectively. The efficiency of ethanol conversion was 0.39g^e^/g^g^, and the total convertibility was 0.45g^e^/g^g^, which represents 90% of the theoretical conversion regardless of the utilized glycerol in cell formation (Table 5).

**Table 5.** Fermentation characteristics of glycerol as a sole carbon source by the generated strain SK-FGG at different oxygen availabilities.

| Parameter name | Fermentation conditions and products values |  |  |  |
| --- | --- | --- | --- | --- |
| Oxygen availability# | Strict anaerobic | Micro-aerobic | Semi-aerobic | Aerobic |
| Fermentation time (h) | 96 ± 0.30 | 60 ± 0.30 | 48 ± 0.30 | 36 ± 0.30 |
| Total consumed glycerol (g/L) | 0.00 ± 0.45 | 37.17 ± 2.66 | 54.01 ± 4.88 | 46.45 ± 0.74 |
| Rate of consumption (g/L<sup>-1</sup>h<sup>-1</sup>) | 0.0 ± 0.0 | 0.62 ± 0.4 | 1.12 ± 0.037 | 1.29 ± 0.035 |
| Total ethanol produced (g/L) | 0.0 ± 0.0 | 15.71 ± 1.87 | 20.97 ± 1.25 | 17.97 ± 0.53 |
| Rate of ethanol production (g/L<sup>-1</sup>h<sup>-1</sup>) | 0.0 ± 0.0 | 0.25 ± 0.04 | 0.44 ± 0.02 | 0.5 ± 0.01 |
| Efficiency of ethanol production (g<sup>e</sup>/ g<sup>s</sup>) | 0.0 ± 0.0 | 0.42 ± 0.05 | 0.39 ± 0.01 | 0.39 ± 0.01 |
| Efficient/ theoretical (%) | 0 | 84 | 78 | 78 |
| Acetic acid accumulation (g/L) | 0.0 ± 0.0 | 0.87 ± 0.21 | 2.88 ± 0.19 | 3.04 ± 0.09 |
| Total conversions (g<sup>p</sup>/ g<sup>s</sup>) | 0.0 ± 0.0 | 0.45 ± 0.04 | 0.44 ± 0.01 | 0.45 ± 0.01 |
| Cell density (OD<sub>600</sub>) | 4.66 ± 0.1 | 8.55 ± 0.27 | 8.55 ± 0.1 | 9 ± 0.1 |

#; Oxygen availability in (liquid medium: flask volume) at 30°C with shaking at 200 rpm with initial glycerol concentration 69 ± 0.7 g/L: Micro (20:100); Semi, (20:200); Aerobic, (20:300).

## DISCUSSION

### The ancestral capability to grow on glycerol

The ancestor strain showed no growth (a maximum OD_600_ after 180h was 0.28) on glycerol as a sole carbon provided in YNB medium (without amino acids) supplemented by 20 mg/L of leucine, histidine, and 5 mg/L of uracil. One of the parent ancestors (25, 29, 30) belongs to S288C. Swinnen et al. reported that the S288C strain is classifying as a negative glycerol grower on a synthetic medium without supporting supplements (14). Hence, herein we initially focused on using a conventional YP medium to avoid these limitations.

### Comprehend the modeling of glycerol fluxes to bioethanol in *S. cerevisiae*

Introducing an active *OpGDH* gene is the first key for deciphering glycerol fermentation, where the activity of endogenous glycerol dehydrogenases was not observed (Table 3). Overexpressing *ScGCY1* alone or combined with the whole DHA is much lower than *OpGDH* here (Data not shown). Although the sole overexpresses of *OpGDH* was not enough to induce efficient conversions (Fig. 3), where the raises of NADH/NAD^+^ ratio by 45% (Table 3) presumably promote oxidizing this plethora through the other NADH-dependent shuttles. At this juncture, Heterologous overexpressing a more dynamic shuttle with the reduced circulation of DHAP into glycerol biosynthesis or/and G3P—glycerolipid— was confirmed as a new point for support fermentation of glycerol or/with glucose. Heterologous expression *NoxE* in *S. cerevisiae* has been successfully oxidizing the excess of NADH during glucose fermentation to produce ethanol or 2,3-butanediol or acetoin. (31-35). By integrating *LlNoxE* while replacing *ScGPD1* within the GN strain, it was showed substantial improvement in ethanol production, which reached 28% compared with the GDH strain within the studied conditions (Fig. 3). This effectiveness of replacing *ScGPD1* with *LlNoxE* shuttle was further confirmed by restoring the balance of intracellular NADH/NAD^+^ ratio (Table 3).

Also, DHAP has been reported for distribution partially into phospholipid and methylglyoxal biosynthesis (36, 37). Therefore, further reducing the ramification of DHAP could be overcome with the overexpressing the *ScTPI1*. The pivotal role of *ScTPI1* is also known toward glycerol production (38). Moreover, overexpressing *ScDAK2* with *ScDAK1* and *CuFPS1*(39) supported the fermentation of glycerol to ethanol. Overexpressing *ScDAK2* (which has a much lower *Km*_(DHA-ATP)_) with *ScDAK1* was reported to detoxified the DHA (40). By integrating the genes in GN-FDT strain, continued bioethanol production until the consumption of glycerol is completed. Nonetheless, the ethanol titer was 20.95 g/L represents less than 48% of fermentation efficiency (Fig.3). Owing to enhanced activities of *ScTPI1, ScDAKs, OpGDH*, and enhance the channeling of glycerol, we overcame re-consuming the produced ethanol earlier than the occurrence of consumption of glycerol (Fig. 3). Thus, the increased activities of *ScDAKs* with *ScTPI1* represent another essential step with both *OpGDH* and *CuFPS1* for glycerol conversion to ethanol. Moreover, the lowering of NADH/NAD^+^ ratio (Table 3).

### Overexpress multi-copies of DHA pathway

With scrutinizing the result, we recognized the conversion efficiency may still be affected by the robustness of natively programmed glycolysis toward glycerol biosynthesis. Furthermore, the activity of alcohol dehydrogenase (ADH) within cells grown in glycerol yeast extract medium has been shown to be ten times higher as compared to those cultured in glucose (41). For overcoming this, we investigated the reinforces the influxes of glycerol with another copy of the DHA pathway. Besides, support the availability of ATP for *ScDAKs* through replacing a phosphorylation pathway through *GUT1* (16). Here, with the integration of the second copy of genes *CuFBS1, OpGDH, ScDAK1*, and *ScTPI1* with highly constitutive expressing systems during replacing *ScGUT1* (22, 42, 43), we extend production levels and efficiencies from 0.85-1.28 g/L^-1^h^-1^ and the conversion efficiency reached 98% of the theoretical ratio in the YP medium (Table 4 [A]). Furthermore, the strain overrode the glucose repression during glycerol fermentation with glucose and produced up to 8.6% of ethanol (Table 4 [D]).

### Ferment glycerol as a sole carbon

Interestingly, flipped her nature even from non-growing on glycerol with this systematic metabolic engineering, where SK-FGG strain showed abilities to convert glycerol in YNB medium—without supplementary amino acids. In micro and semi-aerobic conditions, ethanol production rates were 0.25 and 0.44 g/L^-1^h^-1^ with efficiencies of 0.42 and 0.39g^e^/g^g^, and ethanol titers at 15.7 and 20.97 g/L respectively (Table 5). The remarkable difference between YNB and YP media is the increased accumulation of acetic acid toward YNB from 14 to 53 mg acetic / g glycerol in semi-aerobic conditions (Tables 4, 5). Therefore, the total conversions from the YNB medium reached 90% of the theoretical value regardless of glucose utilized for cell propagation (Table 5). Discuss the results presented here with the introductory reported study (17) is incomplete due to the differences in genetic backgrounds, experimental aims, and the comparable data. Further studies are needed to clarify whether competing aldehyde dehydrogenase (ALD) with GDH on NAD^+^ is the cause and which ALD isoform is responsible for this accumulation.

The content of amino acids in the defined medium has a crucial role in cell growth (44); furthermore, we observed biosynthesis of NAD^+^ from tryptophan through the kynurenine pathway (45, 46). Nicotinic acid, nicotinamide, quinolinic acid, and nicotinamide riboside can salvage the NAD^+^ biosynthesis (28). In this regard, nicotinic acid is auxotrophic under anaerobic conditions in *S. cerevisiae* (46). We also observed different growth and fermentation with the supplements of amino acids or the buffers using SK-FGG strain (Data not shown). These limitations may explain these lower production titers, conversion rates, and efficiency when using YNB compared with the YP medium. We must emphasize that strain SK-FGG was unable to ferment glycerol under the strictly anaerobic condition in our experiment, which used YNB medium (Table 5). Undoubtedly, there are no shuttles for the renovation of NADH under these conditions. Hence, recycling NADH/NAD^+^ represents the complement element for the robust oxidation of the DHA pathway and efficient utilizing glycerol to produce bioethanol. We are currently working on further engineering for glycerol fermentation under anaerobic conditions while using the high reduction merit of glycerol for improving the fermentation efficiencies of lignocellulose’s sugars. Besides, determining the amino acids that may play a significant role with defined media.

In summary, we show here the efficient modeling of glycerol traffic to ethanol production in *S. cerevisiae*. This systematic metabolic engineering includes integration of the following: (i) imposing robust expression of all genes in the DHA pathway; (ii) prevalence of glycerol oxidation by an oxygen-dependent dynamic by the water-forming NADH oxidase *LlNoxE*, which controls the reaction stoichiometries with the regeneration of the cofactor NAD^+^; (iii) revoking the first step of both glycerol biosynthesis and glycerol catabolism through G3P. Our study provides innovative addition to metabolic engineering of re-routing glycerol traffic in *S. cerevisiae* while tracking ethanol production to levels that have not yet been attained within any other safe model organisms, either native or genetically engineered (13, 17, 18, 47-49). The metabolic engineering strategy represents another pivotal step for fermenting glycerol in several promising biorefinery scenarios using the glycell process.

## Funding

This work was supported by Mission 5-2 Research Grant from the Research Institute for Sustainable Humanosphere, Kyoto University.

## Author contributions

S.M.R.K. was responsible for the research idea, conception, planning and organisation of the experiments. S.M.R.K. also provided the information for purchasing the strains, chemicals, and toolboxes for the genetic engineering, performed the experiments and analysed and discussed the results. S.M.R.K. wrote, revised, and submitted the manuscript. T.W. was responsible for all financial support and provided all chemicals and equipment. T.W. was involved in the research idea, the conception, the planning and organisation, the discussion of the results and the manuscript revision and submission.

## Competing interests

The authors state that there are no competing interests to declare.

## Notes

### Competing Interest Statement

The authors have declared no competing interest.

