## Supplementary Materials for "Comprehensive metabolic engineering for fermenting glycerol efficiently in *Saccharomyces cerevisiae*"

Table S1: Primers used within this study.

| Section 1- primers for amplify the genes. | |
|---|---|
| DAK1 F | 5'-atgatggaattcatgtccgctaaatcgtttg-3' |
| DAK 1 R | 5'-catcataagcttttacaaggcgctttgaacc-3' |
| DAK2 F | 5'-atgatggaattcatgtctcacaaacaattcaaatc-3' |
| DAK 2 R | 5'-catcatggatccctagtacccagcagctgtaac-3' |
| TPI1 F | 5'-atgatggaattcatggctagaactttctttg-3' |

| | |
|---|---|
| TPI1 R | 5’-catcatgaattcttagtttctagagttgatgatatc-3’ |
| Δ XhoI site DAK2 F | 5’-taaaaaccatacgttgcgcttggaagatgctgctcttggtatcg-3’ |
| Δ XhoI site DAK2 R | 5’-cgataccaagagcagcatcttccaagcgcaacgtatggtttta-3’ |
| Section 2- primers for sequencing the genes | |
| DAK1 Seq 400 | 5’-atgttgcagttggcagagaaaagggtgg-3’ |
| DAK1 Seq 907 | 5’-ttgctggcacattgatgacctccttc-3’ |
| DAK1 Seq 1670F | 5’-gtatgtcggcgattcatctc-3’ |
| DAK2 Seq 401 | 5’-ggttgctgtaggagatgatgtctctgtc-3’ |
| DAK2 Seq 895 | 5’-tgcccaagagaacgattactggagcattc-3’ |
| TPI1 Seq 401 | 5’-gaaggccggtaagactttggatgttg-3’ |
| CjFPS 928f Seq | 5’-ttcggctacgacatcagaaa-3’ |
| Opgdh 533R Seq | 5’-ttcaacagcatgccaggc-3’ |
| Opgdh 495F Seq | 5’-attcccgacgatatcgga-3’ |
| Section 3- primers for construct URA3:TDH3-d22DIT1t plasmids with partial flanking sequences to GPD1p&t. | |
| GPD1p upstream-TDH3p | 5’-aactcgagtgtatattgtacacccccccctccacaaacacaaatattgataatataaagcccgggagctcagttcgagt-3 |
| Not1-TDH3p R | 5’-tcatgcggccgctttgtttgtttatgtgtg-3’ |

| | |
|---|---|
| NotI *Op*GDH F | 5’-ttgcggccgcatgaaaggtttactttatt-3’ |
| BamHI *Op*GDH R | 5’-aaaggatccttaggacacctcgttcggag-3’ |
| BamHI-d22DIT1t F | 5’-aaataaggatcctaaagtaagagcgctaca-3’ |
| Not1-BamHI- d22DIT1t F | 5’-tttttgcggccgcggatcctaaagtaagag-3’ |
| Not1-NoxE L. lactis F | 5’-caaagcggccgcatgaaaatcgtagttat-3’ |
| BamHI-NoxE L. lactis R | 5’-actttaggatccttattttgcatttaaagc-3’ |
| XhoI-d22DIT1t rev | 5’-aaaactcgaggatgaaaagaaaggcaaat-3’ |
| GPD1t downstream | 5’-ttgtcgacgaaaaaagtgggggaaagtatgatatgttatctttctccaataaatccgcgggatgaaaagaaaggcaaata-3 |
| Section 3.1- Primes for preparing homologous repairing cassettes | |
| Extend GPD1p flanking sequences F | 5’-caagaaacaattgtatattgtacaccccccc-3’ |
| Extend GPD1t flanking sequences R | 5’-tagaagagcctcgaaaaaagtgggggaaa-3’ |
| Section 3.2- Primes for preparing multiplex pCAS-gRNA-GPD1plasmids | |
| gRNA- 135-GPD1 F | 5’-ctggggtactactattgccagttttagagctagaaatagc-3’ |
| gRNA- 135-GPD1 R | 5’-tggcaatagtagtaccccagaaagtcccattcgccacccg-3’ |
| gRNA- 1045-GPD1 F | 5’-cacgaatggttggaaacatggttttagagctagaaatagc-3’ |
| gRNA- 1045-GPD1 R | 5’-catgtttccaaccattcgtgaaagtcccattcgccacccg-3’ |

| | |
|---|---|
| pCAS F | 5'-cggaataggaacttcaaagcg-3' |
| pCAS R | 5'-ttttttctgcagcgaggagcc-3' |
| GPD1p integration check | |
| GPD1t integration check | 5'-ccttaacaagaacaatgtcatgacattgga-3' |
| GPD1 F | 5'-atgtctgctgctgctgatag-3' |
| GPD1 R | 5'-ctaatcttcatgtagatctaattcttcaa-3' |
| Section 4- primers for construct pAUR101; PGKp-*Cu*FPS1-RPL41Bt plasmid. | |
| SmaI-PGKp | 5'-aaaaacccggggagcttggaaagatgcc-3' |
| NotI-PGKp R | 5'-ggaaagcggccgcgatcttttggttttata-3' |
| NotI CuFPS F | 5'-ttgcggccgcatgacaggagaattacttg-3' |
| NotI CuFPS R | 5'-aagcggccgcttaagcgtcaagacgaccg-3' |
| NotI -RPL41t F | 5'-tttttgcggccgcggattgagagcaaatcg-3' |
| SacI- RPL41t R | 5'-aaagagCTCAGCCGAAAATCTTTCAAGCA-3' |
| Section 5- Primers for Gibson assembly module M1- flanking GUT1p&t | |
| F1 FOR primer (**GUT1**-TEF1 p) | 5'-ccatataaaatataccatgtggtttgagttgtggccggaactatacaaatagttatatagcttcaaaatgtttctactcc-3' |
| F1 REV primer (**CuFPS**-TEF1p) | 5'-gtaattctcctgtcatttgtaattaaaacttagattagattgctatgctttctttc-3' |

| | |
|---|---|
| F2 FOR primer (TEF1p – **CuPS**) | 5'-ctaagttttaattacaaaatgacaggagaattacttgctagtggtgaag-3' |
| F2 REV primer (CYC1t -**CuFPS**) | 5'- ggaaaaggggcctgttcaagcgtcaagacgaccgtggctagcctccg -3' |
| F3 FOR primer (**CuFPS**-CYC1t) | 5'- cgtcttgacgcttgaacaggccccttttcctttgtcgatatcatg -3' |
| F3 REV primer (**TYS1p**-CYC1t) | 5'-gtaagcgcaaggacaaattaaagccttcgagcgtcccaaaacc-3' |
| F4 FOR primer (CYC1t- TYS1p) | 5'- cgctcgaaggctttaatttgtccttgcgcttactcgaataggcctccctagc-3' |
| F4 REV primer (OpGDH-TYS1p) | 5'- cctttcatgttatcgtcaattagagtatgcggttatggatgc-3' |
| F5 FOR- TYS1p-linked-**OpGDH gene** | 5'-catactctaattgacgataacatgaaaggtttactttta-3' |
| F6 FOR- OpGDH gene-link ATP15t | 5-ctccgaacgaggtgtcctagtttaacgcttcctgggaactgcagctc -3' |
| F6 REV primer (ATP15t-TDH3p) | 5'-gataaactcgaactgagaggctgaaggcagagaagtttctggaac-3' |
| F7 FOR primer (ATP15t-TDH3p) | 5'-cttctctgccttcagcctctcagttcgagtttatcattatcaatactgccatttc-3' |
| F7 REV primer (DAK1-TDH3p) | 5'-cgatttagcggacattttgtttgtttatgtgtgtttattcgaaac-3' |
| F8 FOR primer (TDH3p-DAK1) | 5'-cacacataaacaaacaaaatgtccgctaaatcgtttgaagtcacagatcc-3' |
| F8 REV Primer (d22-DITIt - DAK1) | 5'-gcgctcttactttattacaaggcgctttgaaccccttcaaaaactc-3' |
| F9 FOR DAK1-linked-d22DITIt | 5'-ggggttcaaagcgccttgtaataaagtaagagcgc-3' |
| F10 FOR FBA1p-linked-d22DITIt | 5'- gcctttcttttcatcataacaatactgacagtac-3' |
| F10 REV Primer (TPI1-FBA1p) | 5'-caaagaaagttctagccatttgaatatgtattacttggttatgg-3' |

| | |
|---|---|
| F11 FOR primer (FBA1p-TPI1) | 5'-ccaagtaatacatattcaaaatggctagaactttctttgtcggtggtaac-3' |
| F11 REV Primer (TDH3t-TPI1) | 5'-gatttaaagtaaattcacttagtttctagagttgatgatatcaac-3' |
| F12 FOR primer (TPI1-TDH3t) | 5'-caactctagaaactaagtgaatttactttaaatcttgcatttaaataaattttc-3' |
| F12 REV Primer (GUT1- TDH3t) | 5'-tggagaggaatataaaattatggaaattacattgttaatagaaattatttatgttgaagggaaagatatgagctatacag-3' |
| F6 extend Rev2 | 5'- tattgataatgataaactcgaactgagagg-3' |
| F7 extend For2 | 5'- cagaaacttctctgccttcagcct-3' |
| Sac1- GUT1p M1 For2 | 5'-agcgagctcgaaccatataaaatatacca-3' |
| Sma1-Gut1t M1 Rev2 | 5'-aacccgggtggagaggaatataaaattat-3' |
| Section 6.1- Primes for preparing multiplex pCAS-gRNA-GUT1 plasmids | |
| pCAS-gRNA target 616 GUT1 F | 5'-attctgtggtcccgccgcacgttttagagctagaaatagc-3' |
| pCAS-gRNA target 616 GUT1 R | 5'-gtgcggcgggaccacagaataaagtcccattcgccacccg-3' |
| Section 6.2- Primes for confirming multiplex pCAS-gRNA-GUT1 system | |
| GUT1p integ. Check F | 5'-cggataaggtgtaataaaatgtg-3' |
| F1 REV primer (***Cu*FPS**-TEF1p) | 5'-gtaattctcctgtcatttgtaattaaaacttagattagattgctatgctttctttc-3' |
| GUT1t integ. Check R | 5'-tcttcataatactagtgttacagtc-3' |
| TPI1 Seq 401 | 5'-gaaggccggtaagactttggatgttg-3' |

| | |
|---|---|
| <u>Barcode</u> | 5’-taagatgtcc <u>aaaccctttgggaaaccctttggg aaaccctttgggaaac</u> gcagcgtacg-3’ |
| Upstream barcode flanking FOR | 5’-caaacggtctcccaccttacaaggtaatatgcatgggtatagcaaacatgtaagatgtcc-3’ |
| Downstream barcode flanking REV | 5’-ttttgccatttgtctctaacgattttgatcgttctggtgtcgttccaaaccgtacgctgc-3’ |

Table S2. Full sequences of the integrated cassettes and module M1 with flanking sequences

1- Cassette1; partial end of GPD1promoter-TDH3p-d22DIT1t-partial front side of GPD1terminator

AA**ctcgag**TGTATATTGTACACCCCCCCCCTCCACAAACACAAATATTGATAATATAAAG**cccgggagctc**AGTTCGAGTTTA
TCATTATCAATACTGCCATTTCAAAGAATACGTAAATAATTAATAGTAGTGATTTTCCTAACTTTATTTAGTCAAAAA
ATTAGCCTTTTAATTCTGCTGTAACCCGTACATGCCCAAAATAGGGGGCGGGTTACACAGAATATATAACATCGTAG
GTGTCTGGGTGAACAGTTTATTCCTGGCATCCACTAAATATAATGGAGCCCGCTTTTTAAGCTGGCATCCAGAAAAAA
AAAGAATCCCAGCACCAAAATATTGTTTTCTTCACCAACCATCAGTTCATAGGTCCATTCTCTTAGCGCAACTACAGA
GAACAGGGGCACAAACAGGCAAAAAACGGGCACAACCTCAATGGAGTGATGCAACCTGCCTGGAGTAAATGATGAC
ACAAGGCAATTGACCCACGCATGTATCTATCTCATTTTCTTACACCTTCTATTACCTTCTGCTCTCTCTGATTTGGAAA
AAGCTGAAAAAAAAGGTTGAAACCAGTTCCCTGAAATTATTCCCCTACTTGACTAATAAGTATATAAAGACGGTAGG
TATTGATTGTAATTCTGTAAATCTATTTCTTAAACTTCTTAAATTCTACTTTTATAGTTAGTCTTTTTTTTAGTTTTAAA
ACACCAAGAACTTAGTTTCGAATAAACACACATAAACAAACAAA**gcggccgcggatcc**TAAAGTAAGAGCGCTACATTGGT
CTACCTTTTTCTTTTACTTAAACATTAGTTAGTTCGTTTTCTTTTTCTTTTTTTATGTTTCCCCCCCAAAGTTCTGATTTT
ATAATATTTTATTTCACACAATTCCATTTAACAGAGGGGGAATAGATTCTTTAGCTTAGAAAATTAGTGATCAATATA
TATTTGCCTTTCTTTTCATCccgcggATTTATTGGAGAAAGATAACATATCATACTTTCCCCCACTTTTTTCgtcgacAA

2- Cassette 2; *Op*GDH cassette, TDH3p- *Op*GDH-d22DIT1t.

**ctcgag**TGTATATTGTACACCCCCCCCCTCCACAAACACAAATATTGATAATATAAAG**cccgggagctc**AGTTCGAGTTTATC
ATTATCAATACTGCCATTTCAAAGAATACGTAAATAATTAATAGTAGTGATTTTCCTAACTTTATTTAGTCAAAAAAT
TAGCCTTTTAATTCTGCTGTAACCCGTACATGCCCAAAATAGGGGGCGGGTTACACAGAATATATAACATCGTAGGT
GTCTGGGTGAACAGTTTATTCCTGGCATCCACTAAATATAATGGAGCCCGCTTTTTAAGCTGGCATCCAGAAAAAAA
AAGAATCCCAGCACCAAAATATTGTTTTCTTCACCAACCATCAGTTCATAGGTCCATTCTCTTAGCGCAACTACAGAG
AACAGGGGCACAAACAGGCAAAAAACGGGCACAACCTCAATGGAGTGATGCAACCTGCCTGGAGTAAATGATGACA
CAAGGCAATTGACCCACGCATGTATCTATCTCATTTTCTTACACCTTCTATTACCTTCTGCTCTCTCTGATTTGGAAAA
AGCTGAAAAAAAAGGTTGAAACCAGTTCCCTGAAATTATTCCCCTACTTGACTAATAAGTATATAAAGACGGTAGGT
ATTGATTGTAATTCTGTAAATCTATTTCTTAAACTTCTTAAATTCTACTTTTATAGTTAGTCTTTTTTTTAGTTTTAAAA
CACCAAGAACTTAGTTTCGAATAAACACACATAAACAAACAAA**gcggccgc**ATGAAAGGTTTACTTTATTACGGTACAA
ACGATATTCGCTACTCCGAAACGGTTCCTGAACCGGAGATCAAAAACCCCAACGATGTCAAGATCAAAGTCAGCTAC
TGTGGAATCTGTGGCACAGACCTGAAAGAATTCACATATTCTGGAGGCCCTGTTTTTTTCCCTAAACACGGCACCAAG
GACAAGATCTCGGGATACGAGCTTCCTCTCTGTCCTGGACATGAATTCAGCGGAACAGTGATTGAGGTTGGCTCTGGT
GTCACCAGTGTGAAACCTGGTGACAGGGTCGCAGTTGAAGCTACGTCCCATTGCTCCGACAGATCGCGCTACAAAGA

| |
|---|
| CACGGTCGCCCAGGACCTCGGGCTCTGTATGGCCTGCAAGAGCGGATCTCCAAACTGCTGTGTGTCGCTGAGCTTCTG<br>CGGTTTGGGTGGTGCCAGCGGCGGTTTTGCCGAGTACGTCGTTTACGGTGAGGACCACATGGTCAAGCTTCCAGACTC<br>GATTCCCGACGATATCGGAGCATTGGTTGAGCCTATTGCTGTTGCCTGGCATGCTGTTGAACGCGCTAGATTCCAGCC<br>TGGCCAGACGGCCCTGGTTCTTGGAGGAGGTCCTATCGGCCTTGCCACCATTCTTGCTCTGCAAGGCCACCGTGCCGG<br>CAAAATTGTGTGTTCCGAGCCGGCCTTGATTAGAAGACAGTTTGCAAAGGAACTGGGCGCTGAAGTGTTTGATCCTTC<br>TACATGTGATGACGCAAATGCCGTTCTCAAGGCTATGGTGCCGGAAAACGAAGGATTCCACGCAGCCTTCGACTGCT<br>CTGGAATTCCTCAGACATTCACCACCTCTATTGTCGCCACAGGCCCTTCGGGAATCGCCGTCAACGTGGCCATTTGGG<br>GAGACCACCCAATTGGATTCATGCCAATGTCTCTGACTTACCAAGAGAAATACGCTACCGGCTCCATGTGCTACACC<br>GTCAAGGACTTCCAGGAAGTTGTCGGGGCCTTGGAAGATGGTCTCATATCTTTGGACAAGGCGCGCAAGATGATTAC<br>AGGCAAAGTCCACCTAAGGGACGGAGTCGAGAAGGGCTTTAGACAGCTCATCGAGCACAAGGAAACCAATGTCAAG<br>ATCCTGGTGACTCCGAACGAGGTGTCCTAA**ggatcc**TAAAGTAAGAGCGCTACATTGGTCTACCTTTTTCTTTTACTTAA<br>ACATTAGTTAGTTCGTTTTCTTTTTCTTTTTTTATGTTTCCCCCCCAAAGTTCTGATTTTATAATATTTTATTTCACACAA<br>TTCCATTTAACAGAGGGGGAATAGATTCTTTAGCTTAGAAAATTAGTGATCAATATATATTTGCCTTTCTTTTCATCccg<br>cggATTTATTGGAGAAAGATAACATATCATACTTTCCCCCACTTTTTTCgtcgac |

3- Cassette 3; NoxE *L.lacis* cassette, partial end of GPD1promoter-TDH3p-*Li*NoxE-d22DIT1t-partial front side of GPD1terminator

**ctcgag**TGTATATTGTACACCCCCCCCCTCCACAAACACAAATATTGATAATATAAAG**cccgggagctc**AGTTCGAGTTTATC
ATTATCAATACTGCCATTTCAAAGAATACGTAAATAATTAATAGTAGTGATTTTCCTAACTTTATTTAGTCAAAAAAT
TAGCCTTTTAATTCTGCTGTAACCCGTACATGCCCAAAATAGGGGGCGGGTTACACAGAATATATAACATCGTAGGT
GTCTGGGTGAACAGTTTATTCCTGGCATCCACTAAATATAATGGAGCCCGCTTTTTAAGCTGGCATCCAGAAAAAAA
AAGAATCCCAGCACCAAAATATTGTTTTCTTCACCAACCATCAGTTCATAGGTCCATTCTCTTAGCGCAACTACAGAG
AACAGGGGCACAAACAGGCAAAAAACGGGCACAACCTCAATGGAGTGATGCAACCTGCCTGGAGTAAATGATGACA
CAAGGCAATTGACCCACGCATGTATCTATCTCATTTTCTTACACCTTCTATTACCTTCTGCTCTCTCTGATTTGGAAAA
AGCTGAAAAAAAAGGTTGAAACCAGTTCCCTGAAATTATTCCCCTACTTGACTAATAAGTATATAAAGACGGTAGGT
ATTGATTGTAATTCTGTAAATCTATTTCTTAAACTTCTTAAATTCTACTTTTATAGTTAGTCTTTTTTTTAGTTTTAAAA
CACCAAGAACTTAGTTTCGAATAAACACACATAAACAAACAAA**gcggccgc**ATGAAAATCGTAGTTATCGGTACGAACC
ACGCAGGCATTGCTACAGCAAATACATTAATTGATCGATATCCAGGCCATGAGATTGTTATGATTGACCGTAACAGT
AATATGAGTTACTTGGGGTGTGGGACAGCTATTTGGGTCGGAAGACAAATTGAAAAACCAGATGAGCTGTTTTATGC
CAAAGCAGAAGATTTTGAAAAAAAGGGAGTAAAGATATTAACAGAAACAGAAGTTTCAGAAATTGACTTTACTAAT
AAAATGATTTATGCCAAGTCAAAAACTGGAGAAAAGATTACAGAAAGTTATGATAAACTCGTTCTGGCAACAGGTTC

ACGTCCAATTATTCCTAACTTGCCAGGAAAAGATCTTAAAGGCATTCATTTTTTAAAACTTTTTCAAGAAGGGCAAGC

CATTGACGAAGAGTTTGCTAAGAATGATGTGAAACGGATTGCTGTGATTGGTGCTGGTTATATTGGGACAGAAATTG

CTGAAGCTGCCAAACGTCGTGGAAAAGAAGTCCTACTTTTTGATGCAGAAAGTACTTCACTTGCTTCATATTATGATG

AAGAGTTTGCTAAAGGGATGGATGAAAATCTTGCCCAACATGGAATTGAACTCCATTTTGGGGAATTAGCTCAAGAG

TTTAAGGCAAATGAAAAAGGTCATGTATCACAGATTGTAACTAATAAATCAACTTATGATGTTGACCTCGTTATTAAT

TGTATTGGCTTTACAGCCAATAGTGCATTGGCTGGTGAACATTTAGAAACCTTTAAAAATGGAGCAATCAAAGTGGA

TAAACATCAACAAAGTAGTGACCCAGATGTTTCTGCTGTAGGAGATGTTGCCACAATCTATTCTAATGCTTTACAAGA

CTTCACCTACATTGCCCTTGCCTCAAACGCTGTTCGCTCAGGGATTGTTGCTGGTCATAATATTGGAGGAAAATCAAT

AGAGTCTGTTGGTGTACAAGGTTCTAATGGAATCTCTATTTTTGGTTACAATATGACTTCTACGGGCTTGTCGGTTAA

AGCTGCGAAAAAAATCGGCCTAGAAGTTTCATTTAGTGATTTTGAAGATAAGCAAAAAGCATGGTTCCTTCATGAAA

ATAATGATAGTGTGAAAATTCGTATCGTTTATGAAACAAAAAATCGCAGAATTATTGGTGCTCAACTTGCTAGCAAG

AGTGAAATAATTGCAGGAAATATTAATATGTTTAGTTTAGCTATTCAAGAAAAGAAAACGATTGATGAATTAGCCTT

ACTTGATTTATTCTTCTTACCACACTTCAATAGTCCATATAATTACATGACTGTTGCAGCTTTAAATGCAAAATAA**ggat**

**cc**TAAAGTAAGAGCGCTACATTGGTCTACCTTTTTCTTTTACTTAAACATTAGTTAGTTCGTTTTCTTTTTCTTTTTTTAT

GTTTCCCCCCCAAAGTTCTGATTTTATAATATTTTATTTCACACAATTCCATTTAACAGAGGGGGAATAGATTCTTTAG

CTTAGAAAATTAGTGATCAATATATATTTGCCTTTCTTTTCATCccgcggATTTATTGGAGAAAGATAACATATCATACTT
TCCCCCACTTTTTTCgtcgac

4- Cassette 5; Glycerol facilitator cassette, PGK promoter- *Candida utilis* glycerol facilitator (*Cu*FPS) - RPL41B terminator.

cccgggAAAGATGCCGATTTGGGCGCGAATCCTTTATTTTGGCTTCACCCTCATACTATTATCAGGGCCAGAAAAAGGA
AGTGTTTCCCTCCTTCTTGAATTGATGTTACCCTCATAAAGCACGTGGCCTCTTATCGAGAAAGAAATTACCGTCGCT
CGTGATTTGTTTGCAAAAAGAACAAAACTGAAAAAACCCAGACACGCTCGACTTCCTGTCTTCCTATTGATTGCAGCT
TCCAATTTCGTCACACAACAAGGTCCTAGCGACGGCTCACAGGTTTTGTAACAAGCAATCGAAGGTTCTGGAATGGC
GGGAAAGGGTTTAGTACCACATGCTATGATGCCCACTGTGATCTCCAGAGCAAAGTTCGTTCGATCGTACTGTTACTC
TCTCTCTTTCAAACAGAATTGTCCGAATCGTGTGACAACAACAGCCTGTTCTCACACACTCTTTTCTTCTAACCAAGG
GGGTGGTTTAGTTTAGTAGAACCTCGTGAAACTTACATTTACATATATATAAACTTGCATAAATTGGTCAATGCAAGA
AATACATATTTGGTCTTTTCTAATTCGTAGTTTTTCAAGTTCTTAGATGCTTTCTTTTTCTCTTTTTTACAGATCATCAA
GGAAGTAATTATCTACTTTTTACAACAAATATAAAACCAAAAGATCG**gcggccgc**atgacaggagaattacttgctagtggtgaaggctgtagttc
tgatatagtattaactaactctacagcagcgccttcctctggtcttgagagaagagcaaacattactgaccacatctcttgtgaacatttcactgctctgcaaagattcagatatggattcagagagtac
tttgctgaatttatcggtaccatgatccttgtgatgtttggtgacggtgttgttgcccagtacactctgtccaagggatctgctggtaactatacaaccattgccttttcgtgggccactgccgttttccttg
gttactgctgttctgcgggtatctctggtgctcatttgaaccctgctgttactctttcagctgctactttcagacagttcccatggagaaaggtattgggttacatgtttgcccaaggtcttggtggttacat

cggcgcccttatcgtttacggtacctatatccaatccatcaacaactactctggtgaaggccagagaatcgccgtcggtgacaaatccacaggtggaatcttctgtactttcccacaaccttacttga

acaccaagggtcaggttacatccgagcttgtcaccactgcccttttgcagtttggtattttctccatgactgatcctcacaatgcaccattgggtaacttcttcccattcggattatggatcttgatttatgg

attgggtacctcttttggttaccagaccggttatgccatcaactttgcaagagatttcaccccaagattggctgctttgactgtcggctatggtaccgagatgttcaccgcctactaccactacttctgg

gtgccaatgatcatcccattcattggtgcattgctcggtgctttcatctatgatttcttcatctaccaaggcttggactcgccattgaaccagccaaagttcggctacgacatcagaaagaagaagatc

caggagtttgaattcaaattggagaactacaagcttgacttcaacccggaggctagccacggtcgtcttgacgcttaagcggccgcGCGGATTGAGAGCAAATCGTTAAG

TTCAGGTCAAGTAAAAATTGATTTCGAAAACTAATTTCTCTTATACAATCCTTTGATTGGACCGTCATCCTTTCGAATA

TAAGATTTTGTTAAGAATATTTTAGACAGAGATCTACTTTATATTTAATATCTAGATATTACATAATTTCCTCTCTAAT

AAAATATCATTAATAAAATAAAAATGAAGCGATTTGATTTTGTGTTGTCAACTTAGTTTGCCGCTATGCCTCTTGGGT

AATGCTATTATTGAATCGAAGGGCTTTATTATATTACCCTTTAGCTTATTCTGAGGTTTCTGTGGCGTGCAAAGTGATG

AACCGGGCGGGTTTTAAGGATAAAATCAAAAAGTGAAAAAATGAACGGAAAATGGAATACCTGTGAAATGGAGAAT

GATAATGAATCTTTCTGTCGTGCTTGAAAGATTTTCGG**CTGAGctc**

5- Module M1; *Cu*FPS1, *Op*GDH, *Sc*DAK1, *Sc*TPI1 cassettes with flanking sequences of GUT1 promoter and terminator

CTCGAACCATATAAAATATACCATGTGGTTTGAGTTGTGGCCGGAACTATACAAATAGTTAT**ATAGCTTCAAAATGT**

**TTCTACTCCTTTTTTACTCTTCCAGATTTTCTCGGACTCCGCGCATCGCCGTACCACTTCAAAACACCCAAGCA**

**CAGCATACTAAATTTCCCCTCTTTCTTCCTCTAGGGTGTCGTTAATTACCCGTACTAAAGGTTTGGAAAAGAAA**

**AAAGAGACCGCCTCGTTTCTTTTTCTTCGTCGAAAAAGGCAATAAAAATTTTTATCACGTTTCTTTTTCTTGAA**

**AATTTTTTTTTTGATTTTTTTCTCTTTCGATGACCTCCCATTGATATTTAAGTTAATAAACGGTCTTCAATTTCT**

**CAAGTTTCAGTTTCATTTTTCTTGTTCTATTACAACTTTTTTTACTTCTTGCTCATTAGAAAGAAAGCATAGCAA**

**TCTAATCTAAGTTTTAATTACAAA**atgacaggagaattacttgctagtggtgaaggctgtagttctgatatagtattaactaactctacagcagcgccttcctctggtcttga

gagaagagcaaacattactgaccacatctcttgtgaacatttcactgctctgcaaagattcagatatggattcagagagtactttgctgaatttatcggtaccatgatccttgtgatgtttggtgacggtg

ttgttgcccagtacactctgtccaagggatctgctggtaactatacaaccattgccttttcgtgggccactgccgttttccttggttactgctgttctgcgggtatctctggtgctcatttgaaccctgctgt

tactctttcagctgctactttcagacagttcccatggagaaaggtattgggttacatgtttgcccaaggtcttggtggttacatcggcgcccttatcgtttacggtacctatatccaatccatcaacaact

actctggtgaaggccagagaatcgccgtcggtgacaaatccacaggtggaatcttctgtactttcccacaaccttacttgaacaccaagggtcaggttacatccgagcttgtcaccactgcccttttt

gcagtttggtattttctccatgactgatcctcacaatgcaccattgggtaacttcttcccattcggattatggatcttgatttatggattgggtacctcttttggttaccagaccggttatgccatcaactttg

caagagatttcaccccaagattggctgctttgactgtcggctatggtaccgagatgttcaccgcctactaccactacttctgggtgccaatgatcatcccattcattggtgcattgctcggtgctttcat

ctatgatttcttcatctaccaaggcttggactcgccattgaaccagccaaagttcggctacgacatcagaaagaagaagatccaggagtttgaattcaaattggagaactacaagcttgacttcaac

ccggaggctagccacggtcgtcttgacgcttgaACAGGCCCCTTTTCCTTTGTCGATATCATGTAATTAGTTATGTCACGCTTACATTCAC

GCCCTCCTCCCACATCCGCTCTAACCGAAAAGGAAGGAGTTAGACAACCTGAAGTCTAGGTCCCTATTTATTTTTTTT

AATAGTTATGTTAGTATTAAGAACGTTATTTATATTTCAAATTTTTCTTTTTTTTCTGTACAAACGCGTGTACGCATGT

AACATTATACTGAAAACCTTGCTTGAGAAGGTTTTGGACGCTCGAAGGCTTTAATTTGTCCTTGCGCTTACTCGAAT

AGGCCTCCCTAGCTATTCTTCAACCTTTCGAACCATCCATACTTCTTACTATCATAATTTTTATTTTATCATGGAGGCG

AGAAGGTCCTTATTCGAGCATCACTAAGAACGGAACTCGAACATTTACAAAGTAGAAAAATTTTATGAAAATTAATT

GTTCTTTCTTCAGAATACAAATTAGTCATTGTCAAAAAGAGATTAGCATCCATAACCGCATACTCTAATTGACGATAA

Catgaaaggtttactttattacggtacaaacgatattcgctactccgaaacggttcctgaaccggagatcaaaaaccccaacgatgtcaagatcaaagtcagctactgtggaatctgtggcacaga

cctgaaagaattcacatattctggaggccctgtttttttccctaaacacggcaccaaggacaagatctcgggatacgagcttcctctctgtcctggacatgaattcagcggaacagtgattgaggttg

gctctggtgtcaccagtgtgaaacctggtgacagggtcgcagttgaagctacgtcccattgctccgacagatcgcgctacaaagacacggtcgcccaggacctcgggctctgtatggcctgcaa

gagcggatctccaaactgctgtgtgtcgctgagcttctgcggtttgggtggtgccagcggcggttttgccgagtacgtcgtttacggtgaggaccacatggtcaagcttccagactcgattcccga

cgatatcggagcattggttgagcctattgctgttgcctggcatgctgttgaacgcgctagattccagcctggccagacggccctggttcttggaggaggtcctatcggccttgccaccattcttgctc

tgcaaggccaccgtgccggcaaaattgtgtgttccgagccggccttgattagaagacagtttgcaaaggaactgggcgctgaagtgtttgatccttctacatgtgatgacgcaaatgccgttctcaa

ggctatggtgccggaaaacgaaggattccacgcagccttcgactgctctggaattcctcagacattcaccacctctattgtcgccacaggcccttcgggaatcgccgtcaacgtggccatttggg

gagaccacccaattggattcatgccaatgtctctgacttaccaagagaaatacgctaccggctccatgtgctacaccgtcaaggacttccaggaagttgtcggggccttggaagatggtctcatat

ctttggacaaggcgcgcaagatgattacaggcaaagtccacctaagggacggagtcgagaagggctttagacagctcatcgagcacaaggaaaccaatgtcaagatcctggtgactccgaac

gaggtgtcctagTTTAACGCTTCCTGGGAACTGCAGCTCTTTTTTTACTCGCTGATATACATTTTAAATATTCTAGCAACTGTG

TATGAAAACTTACGTACTTTTATACGGGAAACTAATAATGACTACAATGATATTGAATACTGGCCGCTTCGAAGAGT

GGTATAAAGTTTGTATCATTGCATTAAAAGAAAAAGAAATATATGTCCCATCATCGCCAATCGCAATGTTGAATGGT

CGTTTACCACTTTTGCGGCTGGGCATATGCAGAAACATGCTGTCCCGTCCCCGACTGGCTAAACTGCCATCTATAAGG

TTTCGGTCTTTGGTCACCCCTTCTTCATCGCAGCTCATTCCTCTCAGTCGGTTGTGTTTAAGGTCACCTGCAGTTGGAA
AATCACTAATTTTACAAAGTTTTAGATGTAATTCATCCAAAACAGTTCCAGAAACTTCTCTGCCTTCAGCCTCTCAGTT
CGAGTTTATCATTATCAATACTGCCATTTCAAAGAATACGTAAATAATTAATAGTAGTGATTTTCCTAACTTTATTTAG
TCAAAAAATTAGCCTTTTAATTCTGCTGTAACCCGTACATGCCCAAAATAGGGGGCGGGTTACACAGAATATATAAC
ATCGTAGGTGTCTGGGTGAACAGTTTATTCCTGGCATCCACTAAATATAATGGAGCCCGCTTTTTAAGCTGGCATCCA
GAAAAAAAAAGAATCCCAGCACCAAAATATTGTTTTCTTCACCAACCATCAGTTCATAGGTCCATTCTCTTAGCGCAA
CTACAGAGAACAGGGGCACAAACAGGCAAAAAACGGGCACAACCTCAATGGAGTGATGCAACCTGCCTGGAGTAAA
TGATGACACAAGGCAATTGACCCACGCATGTATCTATCTCATTTTCTTACACCTTCTATTACCTTCTGCTCTCTCTGAT
TTGGAAAAAGCTGAAAAAAAAAGGTTGAAACCAGTTCCCTGAAATTATTCCCCTACTTGACTAATAAGTATATAAAGA
CGGTAGGTATTGATTGTAATTCTGTAAATCTATTTCTTAAACTTCTTAAATTCTACTTTTATAGTTAGTCTTTTTTTTAG
TTTTAAAACACCAAGAACTTAGTTTCGAATAAACACACATAAACAAACAAAatgtccgctaaatcgtttgaagtcacagatccagtcaattcaag
tctcaaagggtttgcccttgctaacccctccattacgctggtccctgaagaaaaaattctcttcagaaagaccgattccgacaagatcgcattaatttctggtggtggtagtggacatgaacctacac
acgccggtttcattggtaagggtatgttgagtggcgccgtggttggcgaaatttttgcatccccttcaacaaaacagattttaaatgcaatccgtttagtcaatgaaaatgcgtctggcgttttattgatt
gtgaagaactacacaggtgatgttttgcattttggtctgtccgctgagagagcaagagccttgggtattaactgccgcgttgctgtcataggtgatgatgttgcagttggcagagaaaagggtggtat
ggttggtagaagagcattggcaggtaccgttttggttcataagattgtaggtgccttcgcagaagaatattctagtaagtatggcttagacggtacagctaaagtggctaaaattatcaacgacaattt
ggtgaccattggatcttctttagaccattgtaaagttcctggcaggaaattcgaaagtgaattaaacgaaaaacaaatggaattgggtatgggtattcataacgaacctggtgtgaaagttttagaccc

tattccttctaccgaagacttgatctccaagtatatgctaccaaaactattggatccaaacgataaggatagagcttttgtaaagtttgatgaagatgatgaagttgtcttgttagttaacaatctcggcg
gtgtttctaattttgttattagttctatcacttccaaaactacggatttcttaaaggaaaattacaacataaccccggttcaaacaattgctggcacattgatgacctccttcaatggtaatgggttcagtatc
acattactaaacgccactaaggctacaaaggctttgcaatctgattttgaggagatcaaatcagtactagacttgttgaacgcatttacgaacgcaccgggctggccaattgcagattttgaaaagac
ttctgccccatctgttaacgatgacttgttacataatgaagtaacagcaaaggccgtcggtacctatgactttgacaagtttgctgagtggatgaagagtggtgctgaacaagttatcaagagcgaac
cgcacattacggaactagacaatcaagttggtgatggtgattgtggttacactttagtggcaggagttaaaggcatcaccgaaaaccttgacaagctgtcgaaggactcattatctcaggcggttgc
ccaaatttcagatttcattgaaggctcaatgggaggtacttctggtggtttatattctattcttttgtcgggtttttcacacggattaattcaggtttgtaaatcaaaggatgaacccgtcactaaggaaatt
gtggctaagtcactcggaattgcattggatactttatacaaatatacaaaggcaaggaagggatcatccaccatgattgatgctttagaaccattcgttaaagaatttactgcatctaaggatttcaata
aggcggtaaaagctgcagaggaaggtgctaaatccactgctacattcgaggccaaatttggcagagcttcgtatgtcggcgattcatctcaagtagaagatcctggtgcagtaggcctatgtgagt
ttttgaagggggttcaaagcgccttgtaaTAAAGTAAGAGCGCTACATTGGTCTACCTTTTTCTTTTACTTAAACATTAGTTAGTTCGTT
TTCTTTTTCTTTTTTTATGTTTCCCCCCAAAGTTCTGATTTTATAATATTTTATTTCACACAATTCCATTTAACAGAGG
GGGAATAGATTCTTTAGCTTAGAAAATTAGTGATCAATATATATTTGCCTTTCTTTTCATCATAACAATACTGACAGT
ACTAAATAATTGCCTACTTGGCTTCACATACGTTGCATACGTCGATATAGATAATAATGATAATGACAGCAGGATTAT
CGTAATACGTAATAGTTGAAAATCTCAAAAATGTGTGGGTCATTACGTAAATAATGATAGGAATGGGATTCTTCTATT
TTTCCTTTTTCCATTCTAGCAGCCGTCGGGAAAACGTGGCATCCTCTCTTTCGGGCTCAATTGGAGTCACGCTGCCGTG
AGCATCCTCTCTTTCCATATCTAACAACTGAGCACGTAACCAATGGAAAAGCATGAGCTTAGCGTTGCTCCAAAAAA
GTATTGGATGGTTAATACCATTTGTCTGTTCTCTTCTGACTTTGACTCCTCAAAAAAAAAAAATCTACAATCAACAGA

TCGCTTCAATTACGCCCTCACAAAAACTTTTTTCCTTCTTCTTCGCCCACGTTAAATTTTATCCCTCATGTTGTCTAACG
GATTTCTGCACTTGATTTATTATAAAAAGACAAAGACATAATACTTCTCTATCAATTTCAGTTATTGTTCTTCCTTGCG
TTATTCTTCTGTTCTTCTTTTTCTTTTGTCATATATAACCATAACCAAGTAATACATATTCAAAatggctagaactttctttgtcggtggt
aactttaaattaaacggttccaaacaatccattaaggaaattgttgaaagattgaacactgcttctatcccagaaaatgtcgaagttgttatctgtcctccagctacctacttagactactctgtctctttgg
ttaagaagccacaagtcactgtcggtgctcaaaacgcctacttgaaggcttctggtgctttcaccggtgaaaactccgttgaccaaatcaaggatgttggtgctaagtgggttattttgggtcactcc
gaaagaagatcttacttccacgaagatgacaagttcattgctgacaagaccaagttcgctttaggtcaaggtgtcggtgtcatcttgtgtatcggtgaaactttggaagaaaagaaggccggtaaga
ctttggatgttgttgaaagacaattgaacgctgtcttggaagaagttaaggactggactaacgtcgttgtcgcttacgaaccagtctgggccattggtaccggtttggctgctactccagaagatgct
caagatattcacgcttccatcagaaagttcttggcttccaagttgggtgacaaggctgccagcgaattgagaatcttatacggtggttccgctaacggtagcaacgccgttaccttcaaggacaagg
ctgatgtcgatggtttcttggtcggtggtgcttctttgaagccagaatttgttgatatcatcaactctagaaactaaGTGAATTTACTTTAAATCTTGCATTTAAATAAA
TTTTCTTTTTATAGCTTTATGACTTAGTTTCAATTTATATACTATTTTAATGACATTTTCGATTCATTGATTGAAAGCTT
TGTGTTTTTTCTTGATGCGCTATTGCATTGTTCTTGTCTTTTTCGCCACATGTAATATCTGTAGTAGATACCTGATACAT
TGTGGATGCTGAGTGAAATTTTAGTTAATAATGGAGGCGCTCTTAATAATTTTGGGGATATTGGCTTTTTTTTTTAAAG
TTTACAAATGAATTTTTTCCGCCAGGATAACGATTCTGAAGTTACTCTTAGCGTTCCTATCGGTACAGCCATCAAATC
ATGCCTATAAATCATGCCTATATTTGCGTGCAGTCAGTATCATCTACATGAAAAAAACTCCCGCAATTTCTTATAGAA
TACGTTGAAAATTAAATGTACGCGCCAAGATAAGATAACATATATCTAGATGCAGTAATATACACAGATTCCCGCGG

ACGTGGGAAGGAAAAAATTAGATAACAAAATCTGAGTGATATGGAAATTCCGCTGTATAGCTCATATCTTTCCCTTC

AACATAAATAATTTCTATTAACAATGTAATTTCCATAATTTTATATTCCTCTCCACCCGGG

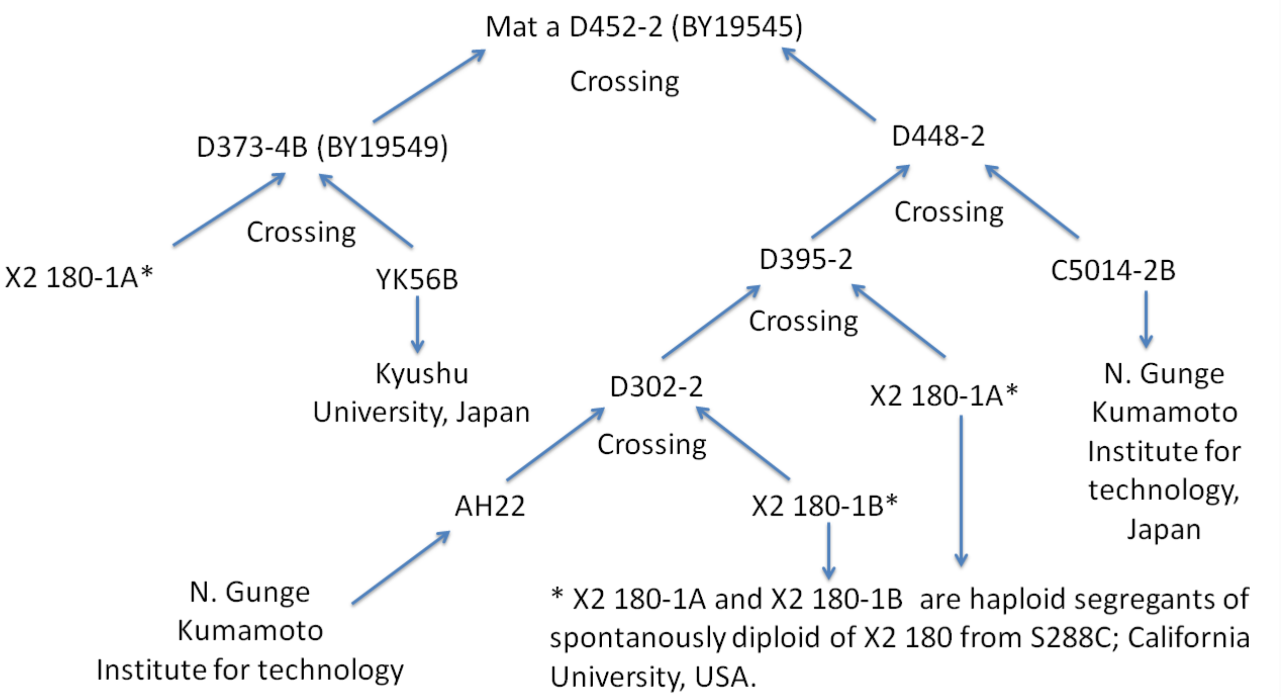

**Fig. S1. Native strain in this study and its ancestors**
